# Genomic and functional gene studies suggest a key role of *beta-carotene oxygenase 1 like (bco1l)* gene in salmon flesh color

**DOI:** 10.1101/821652

**Authors:** Hanna Helgeland, Marte Sodeland, Nina Zoric, Jacob Seilø Torgersen, Fabian Grammes, Johannes von Lintig, Thomas Moen, Sissel Kjøglum, Sigbjørn Lien, Dag Inge Våge

## Abstract

Red coloration of muscle tissue (flesh) is a unique trait in several salmonid genera, including Atlantic salmon. The color results from dietary carotenoids deposited in the flesh, whereas the color intensity is affected both by diet and genetic components. Herein we report on a genome-wide association study (GWAS) to identify genetic variation underlying this trait. Two SNPs on ssa26 showed strong associations to the flesh color in salmon. Two genes known to be involved in carotenoid metabolism were located in this QTL-region: beta-carotene oxygenase 1 (*bco1*) and beta-carotene oxygenase 1 like (*bco1l*). To determine whether flesh color variation is caused by one, or both, of these genes, several functional studies were carried out including mRNA and protein expression in fish with red and pale flesh color. The catalytic abilities of these two genes were also tested with different carotenoids. Our results suggest *bco1l* to be the most likely gene to explain the flesh color variation observed in this population.

## Introduction

All salmon, trout and char species in the genera *Salmo, Parahucho, Oncorhynchus*, *and Salvelinus*, have a unique and characteristic red flesh color ^1^. The color is caused by the binding of carotenoid pigments to muscle alpha-actinin ^2^, and the color intensity is determined both by diet and genetics ^3–5^. The most common muscle carotenoid pigments are astaxanthin and canthaxanthin. Dietary carotenoids are transported over the intestinal brush border, after which they can be metabolized within the enterocytes or in the liver. The portion that remains unmetabolized can be incorporated into lipoproteins, transported into the bloodstream and finally deposited in the muscle ^6^.

Variation in carotenoid metabolism has been linked to the function of *beta-carotene 15,15’-oxygenase* (BCO1) in several species. In humans, genetic variation in both the coding region and the upstream regulatory elements of the *BCO1*, attribute to decreased catalytic activity and increased circulatory levels of carotenes, respectively ^7–10^. In chicken, two SNPs located within the promoter of the BCO1 are associated with breast meat lutein and zeaxanthin content, and hence meat color ^11^. The accumulation of beta-carotene in several mouse organs has been observed following the *BCO1* knockout ^12^.

In mammals, BCO1 provide retinal (vitamin A) through the oxidative cleavage of beta-carotene, a carotenoid which contain two beta-ionone rings. However, in drosophila, chickens, mice, rat, and humans, carotenoids with a single unsubstituted beta-ionone ring can also be cleaved by BCO1 ^13–15^, although the cleavage process is less efficient than for beta-carotene ^16,17^. Carotenoids without any unsubstituted beta-ionone ring, such as astaxanthin and zeaxanthin, cannot be cleaved by BCO1 and are atypical vitamin A sources in mammals ^12,13,18–22^. Interestingly, astaxanthin is the main source of vitamin A in fish ^23–27^, but the molecular pathway for its transformation remains unknown.

In this study we used a whole genome association study (GWAS) and genome sequencing to investigate the genetics underlying the salmon flesh color. The strongest associated SNPs were located within and between two gene paralogs *bco1* and *bco1l* at chromosome 26. These two genes were functionally tested, including the quantification of mRNA and protein expression in red- and pale-fleshed fish. The substrate specificities of salmon’s Bco1 and Bco1l were tested in an *E. coli* strain that was specifically engineered to synthesize beta-carotene and zeaxanthin. The subcellular distributions of Bco1 in the intestinal tissue of the red- and pale-fleshed fish were assessed by the intestine immunostaining and confocal imaging. Finally, RNAseq was used to compare the gene expression profiles in intestinal tissue from red and pale fish, and to investigate the expression of the *bco1* splice variants.

## Results

### Mapping polymorphisms affecting flesh color in Salmo salar

A genome-wide association study including 5650 single-nucleotide polymorphisms (SNPs) identified a strong association between flesh color and a SNP on *Salmo salar* chromosome 26 (ssa26) (**Figure** 1A). The highest scoring SNP in the genome-wide study was positioned in the region encompassing the two paralogous genes *bco1* and *bco1l*. The paralogs were located within the genomic intervals 19.051-19.058Mb and 19.081-19.085Mb, respectively. Considering the already known role of BCO1 in carotenoid metabolism, the two paralogs arose as obvious candidate genes for salmon flesh color.

**Figure 1:**
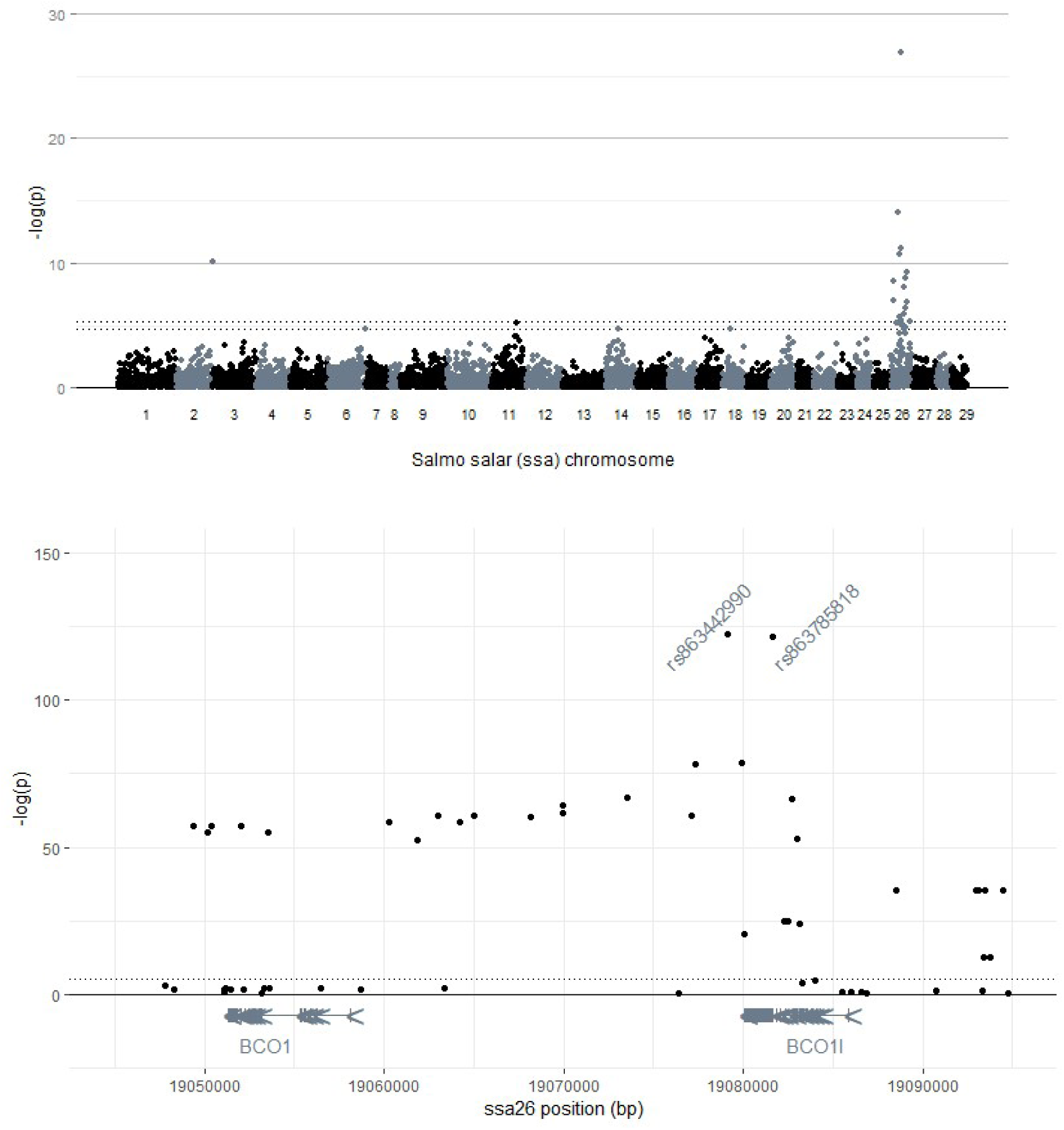
Association studies. A genome-wide association study **(A)** and fine-mapping **(B)** revealed variation within and downstream of the 5’UTR of the *bco1 like* gene (*bco1l*) on *Salmo salar* (Atlantic salmon) chromosome 26 affecting flesh color. Bonferroni adjusted p-value thresholds for α of 0.05 (a) and 0.01 (a and b) are indicated by dotted lines. Positions of the introns and exons, as well as direction of transcription, are indicated for the genes *bco1* and *bco1l*.

Additional fine-mapping revealed the strongest associations (-log(p)>120) with flesh color (Figure 1B) for two SNPs, which are completely linked with each other (r2=0.97): one within (rs863785818), and the other downstream of the 3’UTR of *bco1l* (rs863442990). SNPs with p-values below 1.0e (-log(p)≥ 50) were restricted to the genomic interval of 19.062 to 19.083 Mb of ssa26. No other genes, potentially involved in carotenoid metabolism, were revealed within the given interval. P-values for all the SNPs included in the fine-mapping are given in table S1.

### *bco1* and *bco1l* homeologs in *salmo salar*

Whole-genome sequence data was utilized to identify all *bco1* homeologs resulting from the salmonid-specific WGD (4R) on other *Salmo salar* chromosomes. Homeologs of both *bco1* (GeneID:106562096) and *bco1l* (GeneID:106562095) were found on chromosome 11 (ssa11), positioned in the genomic interval of 18.689 to 18.756Mb.

With respect to *bco1*, ssa26 *bco1* contained 12 exons (GeneID:101448034) while the ssa11 variant contained 11 exons (GeneID:106562096). A stretch of 27 Cytosine’s was inserted 34 bp upstream of exon 2 in ssa11 *bco1*, and a deletion of 37 bp was found in exon 2. The latter introduced a frameshift and a stop codon, making ssa11 *bco1* a pseudogene. The *bco1l* homeologs located on ssa26 (GeneID:101448035) and ssa11 (GeneID:106562095) contain 11 and 10 exons, respectively. A striking difference between these homeologs is a large insert in the intron 1 of ssa11 *bco1l*, which harbors the polycystic kidney disease protein 1-like gene (GeneID:106562168). On ssa11, the coding region is disrupted by a premature stop codon in exon 6, in position 1279 (GeneID:106562095), resulting in a truncated putative protein of 214 amino acids (aa).

A semi-quantitative PCR test revealed no expression of either *bco1* or *bco1l* on ssa11, when using liver cDNA as template. With a genomic DNA as template, PCR amplification produced fragments of the anticipated length at both, ssa26 and ssa11, for *bco1* and *bco1l*. According to our real-time PCR data, the two ssa26 paralogs were mainly expressed in intestine and liver, whereas in both of these tissues *bco1l* was more abundantly expressed than *bco1* (data not shown). These results are also in agreement with the study of ^28^.

### Gene expression differences in red vs. pale fish

A real time PCR assay was used to investigate the gene expression difference between individuals homozygous for rs863785818-allele (T) associated with red muscle and individuals homozygous for the pale muscle associated allele (G) (Figure 2A). When comparing the gene expression of *bco1* in fish with the red- and the pale-associated genotypes, both liver and intestine from fish with the red muscle associated genotype expressed lower levels of *bco1* (p<0.05). *Bco1l* was significantly (p<0.05) more abundant (~3-fold) in the intestine from fish with the red associated genotype, compared to the pale one. In liver, a similar but not significantly different *bco1l* expression pattern was observed (Figure 2A).

**Figure 2:**
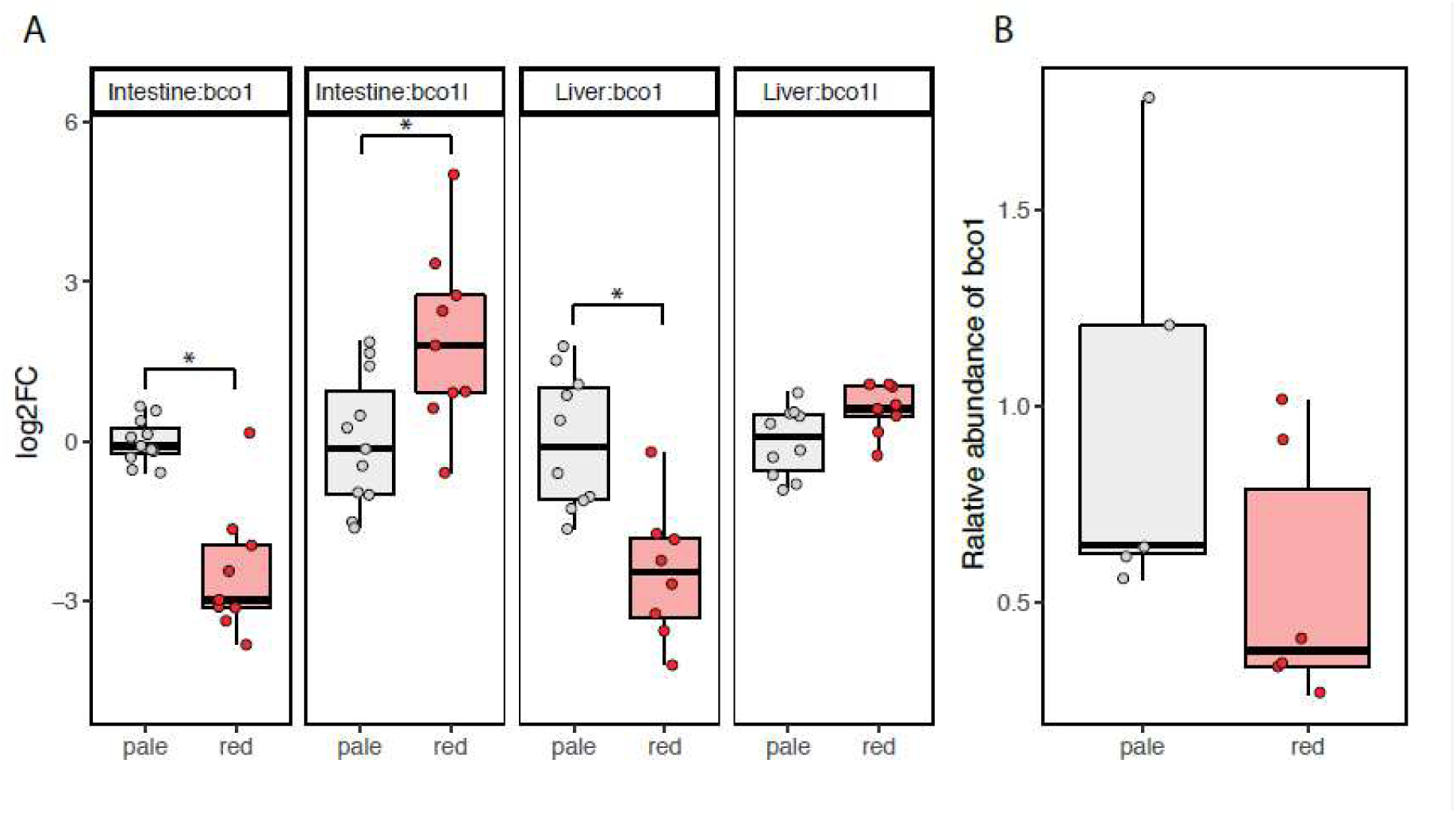
mRNA and protein expression of Bco1 and Bco1 like. **A:** Gene expression of Bco1 and Bco1 like mRNA in liver and muscle of 5 Atlantic salmon being homozygous for rs863785818-allele (T) associated with a red muscle color and 5 individuals homozygous for the pale muscle associated allele (G). Expression is displayed as log_2_ fold changes (logFC) relative to the mean transcription of the gene in the pale group of the respective tissue. An asterisk indicates significantly different expression between the pale and red group (t.test; p.value < 0.05). **B:** Protein expression (quantified Western blots) of Bco1 in Atlantic salmon intestine with red and pale associated genotypes (rs863785818). Protein expression is shown relative to the mean protein expression in the pale group.

### Western blot analysis

Western blot analysis of salmon intestine, using the mouse anti-salmon Bco1 IgG polyclonal antibody, revealed a single band at ~60-kDa, corresponding to the predicted molecular size of Bco1 (Figure 3A). The Bco1 specificity was verified using a peptide competition assay, which showed no signal when the same antibody was pre-incubated with Bco1 blocking peptide in an otherwise identical set-up (Figure 3B). Consequently, the immunoblotting provided solid evidence for the presence of Bco1 protein in the intestine of Atlantic salmon. Bco1 was also two-folds more abundant in the intestine of fish with the pale flesh-associated genotype, compared to those with the red flesh associated genotype (rs863785818 SNP) (Figure 2B). This is also in accordance with the gene expression pattern of *bco1* (Figure 2A). The protein expression pattern of Bco1l could not be measured due to the lack of a specific antibody. Seven antibodies obtained against various epitopes specific to Bco1l were tested. None specifically detected Bco1l, despite extensive protocol adjustments such as different lysis buffers, blocking agents, dilutions, incubation times and temperatures.

**Figure 3.**
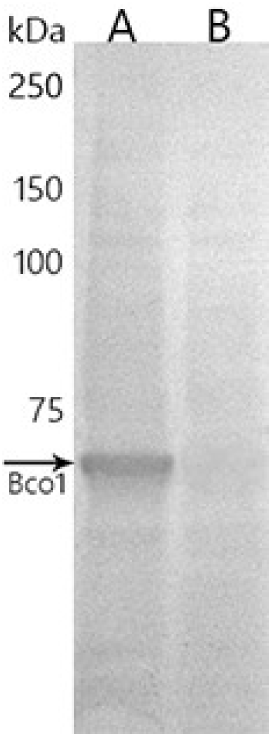
Verification of antibody for Bco1. Western blot analysis of total protein lysate from Atlantic salmon intestine using **(A)** mouse anti-salmon Bco1 IgG polyclonal antibody or **(B)** the same antibody preincubated with the corresponding blocking peptide. Absence of the band in B indicates antibody specificity in the Atlantic salmon intestine. Blots were imaged on a Bio-Rad, ChemiDoc XR and cropped for publication.

### Distribution of Bco1 in Atlantic salmon intestine

Immunofluorescence with the same antibody was used to assess the subcellular distribution of Bco1 in intestine; revealing that the Bco1 is a cytosolic enzyme, most abundant in the apical area of the enterocytes. Bco1 was also present in goblet cells and to a lesser extent throughout the basolateral area of the intestinal villi (Figure 4A). Immunofluorescence signal for Bco1 was markedly more abundant in pale than in red fish (Figure 4A and B), which is in accordance with the western blotting result (Figure 2B).

**Figure 4.**
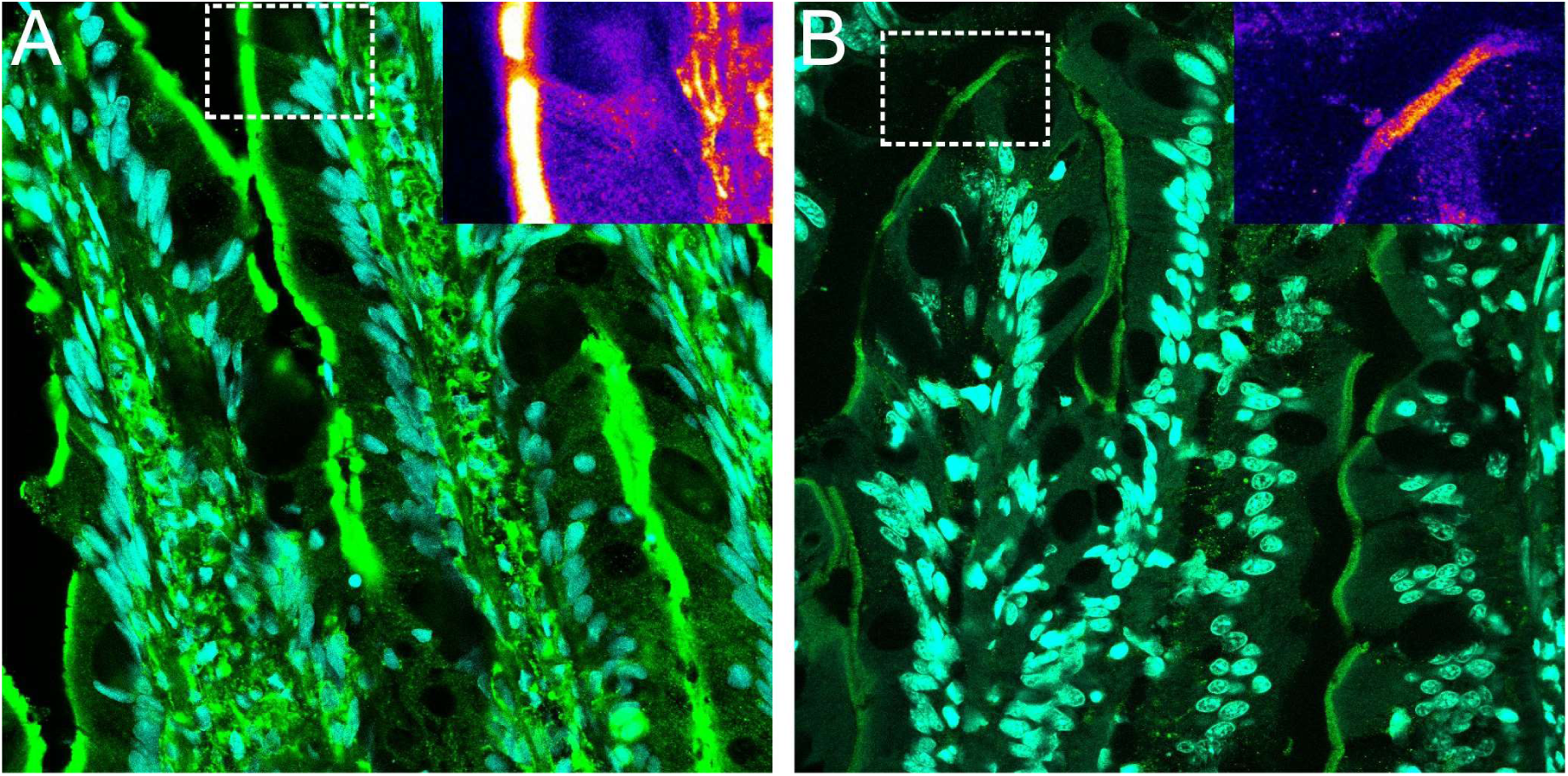
Subcellular localization of Bco1 in Atlantic salmon intestine. Immunofluorescence analysis of Bco1 activity in A. salmon intestines, prepared by freeze substitution and paraffin sectioning (10 μm). Microphotographs were captured on a Leica CLSM SP5, using identical settings, and processed in ImageJ. Bco1 are shown in green and nuclear staining with DAPI in magenta. A) Intestinal Bco1 in pale-fleshed and B) Intestinal Bco1 in red-fleshed Atlantic salmon. Note increased immunofluorescence signal in the pale phenotype. The apical localization of Bco1 in the enterocytes and the abundant activity in goblet cells is common to both phenotypes. Insert images show differences in Bco1 activity corresponding to Fire LUT intensity from ImageJ.

### RNAsequencing

We used RNA sequencing to compare gene expression in the intestine of red-fleshed and pale-fleshed fish (N=11). Among the top ten differentially expressed genes, neither *bco1* nor *bco1l* were detected. Notably, the expression pattern of *bco1* was reversed compared to the real time PCR data. However, the expression difference was not significant when accounting for multiple testing. Gene set enrichment analysis (GSEA) was run on the same dataset of the red vs. pale-fleshed fish, showing significant enrichment of lipid-metabolism related genes in the red-fleshed fish (p< 0.05).

The RNA sequencing dataset was also used to inspect which of the *bco1* variants are present in the salmon intestine. According to the NCBI database, five different *bco1* splice variants are known to exist: XM_014175230, XM_014175231, XM_014175233, XM_014175234 and NM_001279071; with predicted protein lengths of 527, 514, 521, 526, and 522 aa. By counting the reads corresponding to different exon parts, as well as the reads on the exon-exon borders ^29^, we were able to compare the abundance of the various variants. NM_001279071 was detected as the highest expressed *bco1* splice variant in the salmon intestine; based on the high expression of exon 15 along with low expression of exon 9, together with the detected exon spanning reads (Figure 5). Assuming that the most abundant variant also is functional, NM_001279071 was selected for gene and recombinant protein synthesis, as well as for testing of function.

**Figure 5.**
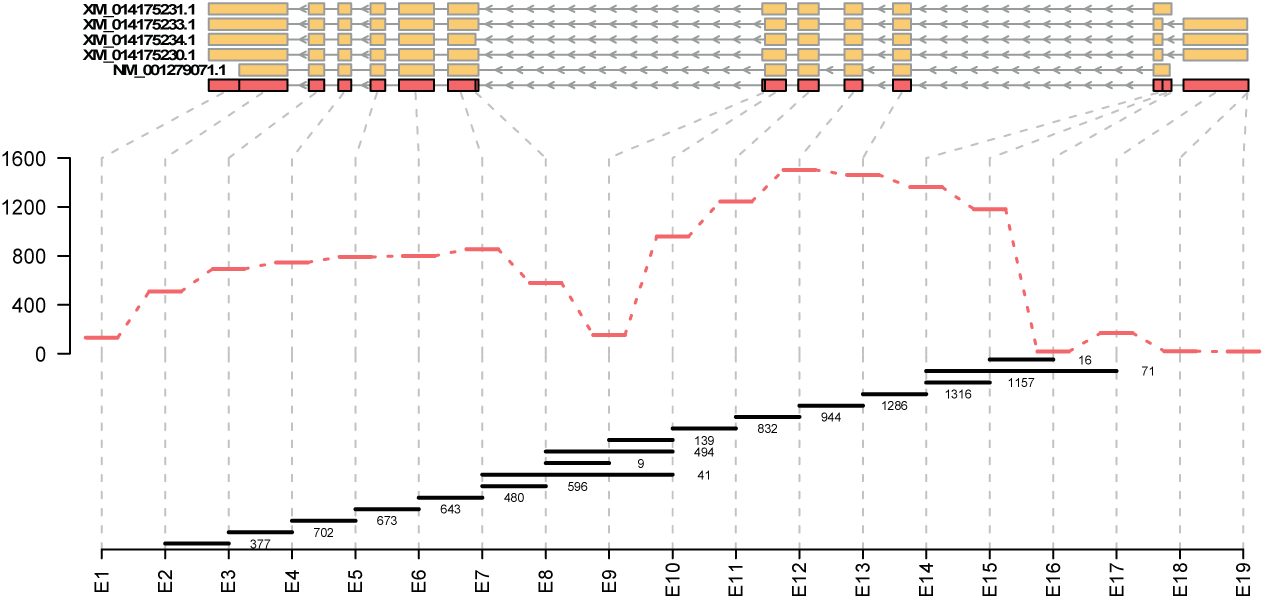
Quantification of alternative bco1 splicing variants. Transcript plot indicating exon usage for *bco1*. The exon structure of the five different *bco1* transcripts is shown in orange. The 19 exon parts derived from the overlap of all five transcripts (representing the counting bins) are shown in red. Read counts for the different exon parts are shown as red bars. The black bars indicate the reads spanning from one exon to another. Start and end point of the horizontal lines indicate which exons are joined, the number to the right shows the exon-spanning read counts.

### Cell based carotenoid cleavage assay

Substrate specificities of the recombinant Bco1 and Bco1l were further tested *in vivo*, using an *E. coli* strain specifically engineered to synthesize beta-carotene or zeaxanthin ^20,30–34^. *Bco1* or *bco1l* expression vectors were introduced to such modified *E. coli*, and the simultaneous expression of *bco1/bco1l* and beta-carotene/zeaxanthin was induced. HPLC was used to detect pigment cleavage products, to measure the enzymes’ ability to cleave the pigments as previously described ^20,30–33^. Following the induction of gene expression, the enzyme activity could also be inferred from the *E. coli* pellet color, which would remain yellow in the absence of enzyme activity. Alternatively, Bco1/Bco1l ability to cleave the pigments would give a color shift from vibrant-yellow to pale-yellow.

The results showed that Bco1l cleaved beta-carotene (Figure 6A). The HPLC chromatograms revealed the formation of retinyl-ester, all-trans-retinal oxime (syn and anti), and all-trans-retinol at the expense of beta-carotene clearly pointing that Bco1l is a 15, 15’ carotenoid oxygenase. Cleavage products observed after expressing *bco1l* in zeaxanthin synthesizing *E. coli* stem from the cleavage of beta-carotene and lycopene (Figure 6B), which are the precursors of zeaxanthin in the pathway. So the two experiments proved that the Atlantic salmon Bco1l has 15, 15’ oxygenase activity, while our efforts failed to express BCO1 in this test system

**Figure 6.**
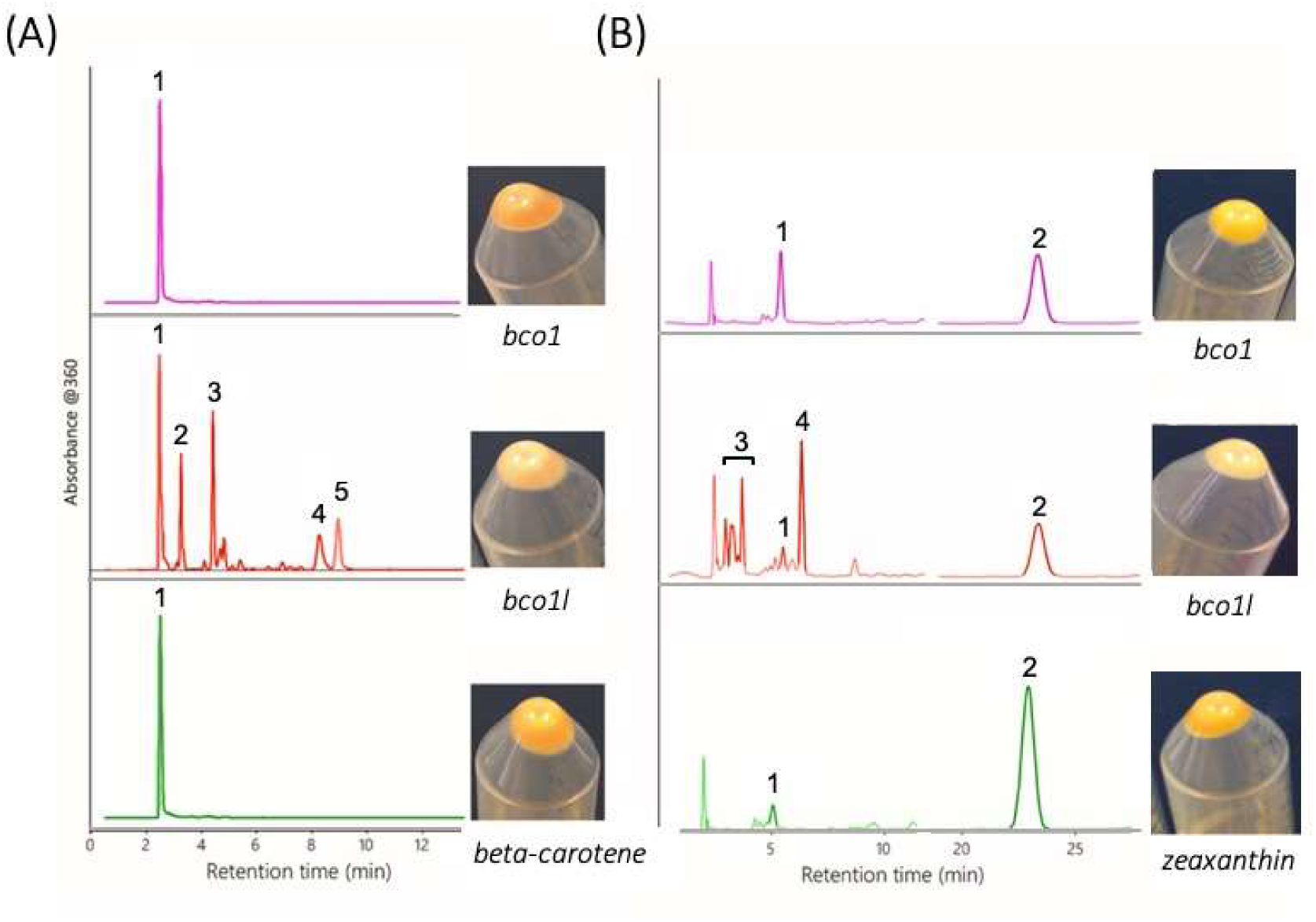
*In vivo* testing of Bco1 and Bco1l in pigment producing strains of *E. coli*. **(A)** An *E. coli* strain specifically engineered to produce beta-carotene were transfected with plasmids carrying either *bco1* or *bco1l* coding sequences. The cellular pellets were analyzed on HPLC and the resulting cleavage products were detected. The HPLC chromatogram in purple shows that Bco1 did not cleave beta-carotene. HPLC chromatogram in red indicates that Bco1l was capable of cleaving beta-carotene into retinol (vitamin A) and its isoforms, proving that Bco1l has 15,15’-oxygenase activity. The HPLC in green shows the control beta-carotene producing *E.coli*, which were not transfected with neither *bco1* nor *bco1l* coding sequence. **(B)** Plasmids carrying either *bco1* or *bco1l* coding sequence was introduced into *E. coli* strain specifically engineered to produce zeaxanthin. The HPLC chromatogram in purple shows that Bco1 had no cleavage activity, leaving only intact zeaxanthin. The HPLC chromatogram in red shows the cleavage products observed following the expression of bco1l in zeaxanthin-producing *E.coli* transfected with plasmids carrying *bco1l* coding sequence. The cleavage products could stem from the Bco1l cleavage of the carotenes beta-carotene and lycopene which are precursors of the xanthophyll zeaxanthin in that pathway. The HPLC chromatogram in green shows the control, zeaxanthin-producing *E.coli* that were not transfected with neither *bco1* nor *bco1l* coding sequences. *E. coli* bacteria pellets are shown to the right of the HPLC chromatograms. Bco1l activity could be inferred from the shift in color of the pellet from vibrant yellow (observed in the control *E. coli*) to pale-yellow (observed in the *E. coli* with induced *bco1l* expression). Such shift in the color of *E. coli* pellet was not observed in neither beta-carotene nor zeaxanthin synthesizing *E. coli*, following the *bco1* expression. The scale bars indicate absorbance of 0.01 at 360 nm. Peaks corresponding to major compounds are denoted with numbers (A) 1: beta-carotene, 2: retinyl-ester, 3: All-trans-Retinaloxime-syn., 4: All-trans-Retinaloxime-anti 5: All-trans-Retinol. (B) 1: beta-cryptoxanthin, 2: zeaxanthin, 3: retinaldehyde isomers, 4: unknown.

### Sequence characteristics of two bco1 homologues in Atlantic salmon

The Atlantic salmon *Bco1* encode a 520 aa protein, while *Bco1l* encode a 528 aa protein. All Bco1 proteins, including the salmon orthologs, had four conserved histidines (H172, H237, H308, H514, according to human BCO1 positions) that coordinate the iron required for BCO1 activity ^35,36^. In addition to the four histidines, eight additional residues are proposed to form the active site of the human BCO1 (F51, F93, S139, E140, T141, Y235, F307, W326) ^35,37^. Of these, salmon BCO1 diverged at F93M and F307Y, while salmon BCO1l diverged at T141V and Y236F. Furthermore, the residues P101, C102, I105, F106, K108, L258, T262 and Y264 form a hydrophobic patch at the entrance to the active tunnel in human BCO1, forming a dome. Salmon Bco1 had substitutions at four of these positions: C102G, I105V, F106I and K108R, while Bco1l only diverged at Y264I (Figure S1).

Furthermore, we compared the Bco1 and Bco1l 3D models. A crystal structure of bovine RPE65 was used as a template ^38^ for making a model to predict how Bco1 and Bco1l might interact with their substrates. The similarity to human RPE65 was 38% identities and 57% positives for Bco1 and 41% identities and 60 % positives for Bco1l, indicating that the 3D model was sensible. This was also supported by the Global Model Quality Estimates (GMQE) score of 0.70. The overall structures of the predicted Bco1 and Bco1l were highly comparable (Figure S2). Bco1 and Bcol1 3D models disjoined at the proteins’ periphery, which is outside the enzymes’ active sites (Figure S3). The disjoined regions encompass aa 102-106, 203-227 and 341-353.

## Discussion

A genome-wide association study was conducted to identify polymorphisms influencing flesh color intensity in salmon. SNPs located in a given region of ssa26 showed the strongest association with flesh color (Figure 1A). This region harbors two genes, *bco1* and *bco1l*, both known to be involved in carotenoid metabolism. Re-sequencing and additional genotyping in this region showed that the two SNPs strongest associated with flesh color were located within the 3’-UTR of *bco1l* and immediately downstream of the *bco1l* gene, respectively (Figure 1B). These two SNPs were in almost complete linkage disequilibrium with each other (r2=0.97), so association alone does not resolve which one might be the causal.

Since both Bco1 and Bco1l are oxygenases with an anticipated ability to cleave carotenoids, one would intuitively expect the causative enzyme(s) to be more active in pale fish, since increased carotenoid cleavage would leave less pigment available for muscle deposition. Looking at the expression profiles in Figure 2A and B, this assumption is compatible with the expression pattern of *bco1*, while *bco1l* have an opposite expression pattern, i.e. is more expressed in individuals with deep red muscle color. Based on the expression profiles alone, *bco1* is more likely to be the causative gene. Western blotting also demonstrated that Bco1 is two-fold more abundant in the intestine of pale compared to the red-fleshed fish.

Immunofluorescence staining of Bco1 in sections of salmon intestine tissue indicated that Bco1 is a cytosolic enzyme, distributed across the enterocyte, with the strongest immunoreactivity in subapical region, close to the membranes of the apical surface of absorptive epithelial cells. This distribution is in line with what has been observed for human BCO1 ^39,40^. However, *bco1l* is expressed at a considerable higher level compared to *bco1* ^28^, possibly indicating that the *bco1l* product has a higher catabolic capacity *in vivo* compared to *bco1*. The abundance of Bco1l protein could unfortunately not be assessed due to our failure of generating a salmon specific Bco1l antibody.

*Danio rerio bco1* knockdown larvae showed signs of impaired retinoic acid-dependent developmental processes ^41^, suggesting a similar role of piscine bco1 in vitamin A formation as in mammals, that could not be rescued by *bco1l*. In mammals, the expression of *bco1* is tightly regulated to ensure sufficient, but avoid a toxic, levels of vitamin A. This strong restriction on *bco1* expression make it a less probable candidate for causing the considerable genetic variation of flesh color. To degrade the significant amounts of carotenoids in the salmon diet, the highly expressed *bco1l* might be a more likely candidate gene, which also will leave the vitamin A homeostasis unaffected. However, it remains to explain how the observed expression pattern of *bco1l* fits this theory.

In order to distinguish between the functions of Atlantic salmon *bco1* and *bco1l*, we also tried to produce the enzymes *in vivo* by expressing *bco1/bco1l* in *E. coli* strains specifically engineered to produce beta-carotene or zeaxanthin. According to the NCBI database, five *bco1* (NM_001279071, XM_014175230, XM_014175231, XM_014175233, XM_014175234) and a single *bco1l* isoforms have been reported in Atlantic salmon. The five *bco1* variants were also identified in our RNAseq data. The NM_001279071 *bco1* variant was the most abundantly expressed one and was mainly found in the intestine. Therefore, the NM_001279071 isoform was selected for the *in vivo* testing. The results indicated that Bco1l is capable of cleaving beta-carotene into vitamin A by 15,15’-oxygenase activity (Fig.6A). However, it is not possible to specifically detect 3-OH-retinoids by HPLC. The observed fragments in Fig. 6B can therefore stem from cleavage of beta-carotene or lycopene, which are the precursors of zeaxanthin in this pathway. It is therefore not possible to conclude from this experiment if Bco1l is capable of cleaving oxygen-containing xanthophylles such as zeaxanthin and astaxanthin. Notably, expression of NinaB, an insect Bco1 that cleaves zeaxanthin, in this test system led to retinal but not3-hydroxy-retinal production ^42,43^. These finding clearly indicate that zeaxanthin is not available as substrate for the recombinant enzymes in the heterologous expression bacterial background.

Protein sequence inspection showed that the four key histidine residues required for enzyme activity in orthologues enzymes were conserved in both Bco1 and Bco1l (Figure S1). The most pronounced difference between Bco1 and Bco1l at known functional positions are related to the patch-forming amino acids. Substitution experiments involving these residues have shown to impact enzyme activity, and position 108 has specifically been shown to influence substrate specificity ^37^. One of the disjoining regions in the 3D-model between Bco1 and Bco1l include aa 102-106, which include 3 of the amino acids making up the enzyme patch (C102, I105, F106). This may indicate that the salmon Bco1 and Bco1l diverge in substrate specificities, but that both enzymes are likely to be functional.

It has previously been shown that fish have evolved and maintained two *Bco1* genes prior to the teleost-tetrapod split ^28^, driven by the presence of unique carotenoids in the aquatic environment, and presumably facilitated trough a gene duplication event. Despite the chromosomal proximity, the *Salmo salar bco1* and *bco1l* genes do not appear to be under common regulation, as there is a substantial difference in gene expression in liver, and overall expression level in gut, liver and muscle ^28^. The two paralogs show only 54% protein identity, further indicating extensive neo-functionalization following the gene duplication. Beta-carotene is a dominant carotenoid in the mammalian diet, which is used for conversion into vitamin A via the activity of BCO1. Astaxanthin is the most abundant carotenoid in the salmon diet and serves as a source of vitamin A, while beta-carotene and other carotenoids are present at lower levels. The yet unknown metabolic pathway of astaxanthin conversion into vitamin A in Atlantic salmon could be enabled through the functional diversification of *bco1* and *bco1l*.

The G > T variant in the 3’UTR of *bco1l* in red-flesh salmon may affect the enzyme activity of Bco1l by creating a target site for microRNAs binding, which impair mRNA translation to protein. Reduced protein translation could then be compensated by higher level of transcription in the red-flesh fish. There are known examples of 3’-UTR polymorphisms generating or destroying miRNA binding sites that significantly affect the phenotype, e.g. ^44^.

In summary, this work provide additional insights into the molecular basis of salmon flesh pigmentation. Two SNPs showed strong association to a region of ssa26, harboring the two paralogs *bco1* and *bco1l.* Fish with a pale flesh color had higher *bco1* expression and more abundant cellular Bco1 proteins compared to the fish with red flesh color, whereas the opposite was true for *bco1l*. In zebrafish the knockdown of *bco1* affected vitamin A dependent processes, indicating: 1) that the regulation of *bco1* is under very strict control and 2) that *bco1* in fish and mammals have comparable roles ^41^. It is therefore unlikely that the *bco1* gene, with such tightly regulated expression, would be the key enzyme in degrading the large and variable amounts of astaxanthin in the salmon diet. In that respect, it appears more likely that the abundantly expressed *bco1l* ^28^ might be the major astaxanthin degrading enzyme.

The sequence analysis and structure prediction of Bco1 and Bco1l strengthened the possibility that the two paralogs have similar cleavage mechanisms, but different substrate specificities. Genetic variation detected in the 3’UTR of *bco1l* could then affect the activity of Bco1l, and consequently the level of intact astaxanthin available for muscle deposition. Neither *in vivo* nor *in vitro* assays provided us definite answers about the substrate specificities of *bco1/bco1l*. *In vivo* functional assays confirmed that salmon Bco1l is an active 15,15’-carotenoid oxygenase. The possibility that Bco1l is capable of utilizing xanthophylls such as zeaxanthin and astaxanthin cannot be excluded, but is not confirmed by this work. Further experiments are needed to unequivocally prove the substrate specificities of *bco1* and *bco1l*. Understanding the exact roles of *bco1/bco1l* would be important in deciphering the metabolism of the diverse family of carotenoids in the salmon diet.

## Materials and methods

### Ethics statement

Data and samples used for the GWA-study are routinely collected as a part of the AquaGen breeding program, in accordance with good husbandry practice and the Norwegian law for aquaculture production and fish breeding given in the Aquaculture Act and more specific in “Regulations on the operation of aquaculture facilities” (Aquaculture Operation Regulations). AquaGen are certified with GlobalGap, ISO9001:2015, RSPCA welfare standards for Salmon and have adopted and follows the code for European responsible breeding “CodeEfabar”. Tissue samples used for the gen/protein expression studies and the immunofluorescent microscopy were also collected from fish of the AquaGen breeding population, in accordance with Regulation of Animal Experiments in Norway and EU2010/63. The fish were killed before taking out tissue samples.

### Family material and phenotypes

Founding sires were sampled from four separate subpopulations that hatched in the years 1998-2001, whereas the dams were sampled from two separate subpopulations that hatched in the years 2000-2001. Offspring from full-sib families were reared in separate tanks until individual tagging, and after tagging all fish were held in the same environment until they reached an age of two years. All fish were starved for two weeks before slaughter at a commercial slaughter house. Pieces of flesh were packed in individual plastic bags and stored at 4°C. Flesh pigment content was determined by analyses of near-infra-red reflections (NIR) in fish flesh using the QMonitor system (Q Vision, Asker, Norway). A more detailed description of the rearing and handling of the fish can be found in Bahuaud *et al*. ^45^.

### Genotyping and re-sequencing

Genomic DNA was extracted from fin clips for a total of 2510 phenotyped fish. Initially, the fish were genotyped according to manufacturer’s instructions using an Atlantic salmon iSelect SNP array developed by Center for Integrative Genetics (CIGENE) ^46^ in Norway. Genotype data were analyzed using Genome Studio (Illumina Inc., San Diego, US) and the R-package beadarrayMSV (http://cran.r-project.org/).

Re-sequencing data from 45 Atlantic salmon provided by Aqua Gen AS was used for SNP detection on ssa26. Reads were aligned to the Atlantic salmon reference genome ^47^ with Bowtie2 ^48^. SNP detection was performed with FreeBayes ^49^. In a 1.24Mb sequence covering and flanking the ssa26 *bco1* genes, a total of 579 SNPs were detected. These SNPs were genotyped in the 2510 phenotyped fish with the Sequenome MassARRAY (Agena Bioscience Inc., San Diego, US) technology. The SNPs without a pre-existing rs-number has been submitted to European Variation Archive (EVA) under Project: PRJEB28258.

### Association mapping

Association analyses were performed with the statistical model:

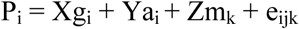

Where P is phenotypic value of individual i, g is fixed effects for individual i, a is random effect of individual i where co-variance structure between individuals is determined from pedigree relationships, m is random effect of genetic marker k, and e is an error term. Fixed effects were gender, weight at slaughter and population of origin for sire and for dam. Estimations were conducted with the ASREML software ^50^. It has been shown by MacLeod *et al.* ^51^ that including effect of individual based on pedigree relationship in the mixed model reduces the number of false positives in genome-wide scans.

The Wald F-statistic and denominator degrees of freedom were estimated with the ASREML software. A Bonferroni correction was applied to significance thresholds.

### Gene expression of bco1 and bco1l homeologs

A real time PCR assay was set up for investigating expression levels of the *bco1* paralogues. A total of 10 fish with and without the QTL was analyzed for expression in the distal intestine and liver. Tissue biopsies were sampled before total RNA extraction (RNeasy mini kit, Qiagen, Hilden, Germany) and cDNA synthesis (QuantiTect revese transcription kit, Qiagen) according to the manufacturers instructions. Real time PCR was optimized and carried out using primers specific to *bco1* (Sense; 5’ AGATCATGAAGTTGGACACTGA, Anti sense; 5’CATCCTCTCCTTCTCCATTG), *bco1l* (Sense; 5’ CTCACTCAATCCAAAGAAATCC, anti-sense; 5’AATGAAGTGTCCATGCAGATC). Efficacy of the PCR assays were >0.93 for all reactions and the data were normalized against *ef1a* (Sense; 5’CTGGACACAGAGACTTTATCAAGA, anti-sense; 5’CTTAGAGATACCAGCCTCAAACTC) and relative expression was calculated using ∆∆Ct.

### Western blotting

Total protein lysates were prepared from intestinal tissue collected from pale and red-fleshed individuals as defined by the genotype of rs863785818 SNP. Five homozygous individuals with the red associated T allele and six homozygous individuals with the pale associated G allele was analysed. This SNP is located within the 3’UTR of bco1l and show, together with rs863442990, the strongest association to flesh color. Protein lysates were prepared from 10 mg tissue using RIPA buffer (Tris-Cl, 50 mM; pH 8.0, 150 mM NaCl, 0.1% SDS, 1% NP-40, 0.5% deoxycholate and 1 mini protease inhibitor cocktail tablets / 50 ml (Roche Applied Science) and centrifuged at 14000g for 30 min at 4°C. Protein concentration of the supernatant was measured using Bradford reagent (Bio-Rad Laboratories), before mixing 10 ug of protein 2x laemmli buffer (63 mM Tris-HCl, pH 6.8, 10% glycerol, 5% 2-mercaptoethanol, 3.5% sodium dodecyl sulfate) and heating at 60°C for 10 minutes. Denatured protein was separated using a 4-15% Mini-PROTEAN® TGX™ Precast Gel (Bio-Rad Laboratories). Proteins were transferred to a PVDF membrane (Bio-Rad) using a Trans Blot Turbo (Bio-Rad) protein transfer system. Membranes were stained with Ponceau S to assess equal loadings and imaged for density measurements in ImageJ (http://imagej.nih.gov/ij/; NIH; Bethesda, MD). After Ponceau red destaining, the blots were blocked with 2% dry milk in TBST (50 mM Tris-HCl, pH 7.4, 200 mM NaCl, 0.1% Tween) overnight at 4°C and five times washed with TBST. Bco1 was detected using a 1:3000 dilution of a polyclonal mouse IgG antibody raised against the Atlantic salmon proteins epitope sequence (KVPDTTAGDKAN) by Abmart (NJ, USA). The blots were incubated with primary antibody in 2% dry milk in TBST overnight at 4°C, before being washed five times in TBST. Signal was detected by incubation with an alkaline phosphatase conjugated secondary goat anti-mouse IgG antibody (ThermoFisher) diluted 1:1000 for 1 hour at room temperature in TBST with 2% dry milk. After five times washing with x TBST and alkaline phosphatase buffer (0.1 M Tris-Cl, 50mM MgCl2, 0.1% Tween 20), the blots were stained with NBT/BCIP solution (Sigma-Aldrich), rinsed with water and dried. Images were captured and density measurements were normalized against the Ponceau red image. To verify the specificity of antibody binding, primary antibody was incubated overnight at 4°C with a 10X molar excess of synthetic epitope peptide. The neutralized primary antibody was then used in side-by-side comparisons with untreated antibody. Samples were imaged on a Bio-Rad, ChemiDoc XR.

### Immunofluorescent microscopy

For immunofluorescent microscopy tissues were sampled from the same fish pool as used for the western blotting protein quantification. Freeze substituted paraplast (Sigma-Aldrich; St Louis, MO) embedded intestine tissues (8 μm in thickness) of red and pale-fleshed Atlantic salmon were used for the immunofluorescence assay. For antigen retrieval, slides were boiled in an autoclave at 95°C in antigen-retrieval buffer (10 mM Tris-base; pH 8.6, 1 mM EDTA, 0.0) for 30 minutes, after which slides in the solution were cooled down to room temperature. The tissue sections were blocked in 4% dry milk powder in 1 x TBST for 30 min followed by overnight incubation at 4°C with anti-Bco1 antibody. The primary antibody was detected by Alexa Fluor® 647 conjugated secondary goat anti-mouse IgG antibody (Thermo Fisher Scientific). Several washes in TBST followed each step. DAPI (Sigma-Aldrich) was used for nuclear counterstaining. Samples were imaged on a Leica TCS LSI-III. Immunofluorescence was evaluated by applying excitation and emission filters adapted for the Alexa Fluor® 647 fluorochromes, recommended by the manufacturer. ImageJ was used for image analysis and processing.

### mRNA sequencing

The fish used for the transcriptome analyses did not originate from the same fish pool as used for the western blotting protein quantification and immunofluorescence, due to the fish shortage. However, the same rs863785818 SNP was used as criteria to select five red and six pale-fleshed individuals for the analysis. Total RNA (RIN> 9.5) was extracted from freeze substituted 300 μm vibratome sliced intestine tissues, using Quiagen RNeasy kit (Qiagen, Germany). Eleven samples were used to produce libraries using an Illumina TruSeq stranded mRNA kit (Illumina, California, USA) manufacturer’s instructions with the following slight modifications: (i) 2µg input RNA, (ii) incubation for 25 s (vs. 8 min) at the “elute, fragment and prime” step, and (iii) using a cDNA:bead ratio of 1:0.7 ratio (vs. 1:1) after adapter ligation and PCR step. Libraries were quantified using a Qubit® 2.0 Fluorometer and the Qubit® dsDNA BR Assay kit (Invitrogen™, Thermo Fisher, Massachusetts) and profiled using a 2100 BioAnalyzer system with the DNA High Sensitivity kit (Agilent Technologies, California), before being diluted to 10 nM and equal volumes being combined into a single pool. Sequencing was performed on a MiSeq machine (Illumina, California) with a v3 reagent kit, generating 2 x 300 paired-end reads.

### Analysis of RNAseq data

RNA sequencing files were processed in the following manner: (i) Illumina adaptors were removed and the sequences were quality-trimmed using cutadapt (v1.8.1 with Python 2.7.8) specifying the options: -q 20 -O 8 --minimum-length 40. (ii) Trimmed reads were mapped to the Atlantic salmon genome (GenBank Accession number: GCA_0002333375.4) using STAR (v2.4.2a) ^52^. (iii) Read alignments, recorded in BAM format were subsequently used to count uniquely mapped reads per gene using the HTseq-count.py script (v0.6.1p1) ^53^, with the RefSeq genome annotation where unique gene_ids were assigned to gene features. All Illumina sequence reads are publicly accessible through the European Nucleotide Archive (http://www.ebi.ac.uk/ena) with the accession PRJEB10297.

Gene expression levels were calculated as counts per million total library counts using the R package edgeR ^54^. Total library sizes were normalized to account for bias in sample composition, using the trimmed mean of m-values approach. Log-fold expression differences were calculated contrasting samples with red and pale flesh color.

Additionally, read alignments (BAM format) were pooled and subsequently processed to count exonic parts for bcmo1 according to the process deciphered in the vignette of the R package DEXSeq ^29^. In short: RefSeq genome annotation was processed using dexseq_prepare_annotation.py script before counting exonic features using the dexseq_count.py script.

### Expression of bco1 and bco1l constructs in carotenoid producing E. coli

Bco1 or bco1l expression vectors were transformed into E. coli XL1blue cells carrying expression plasmids for beta-carotene or zeaxanthin biosynthesis. The transformed E. coli were grown overnight at 37°C on LB solid medium containing 25 µg/mL kanamycin and 35 µg/mL chloramphenicol. Selected colonies were picked into 5 mL LB medium containing 25 µg/mL kanamycin and 35 µg/mL chloramphenicol and grown at 37°C with shaking until the OD600 was ∼0.6. Expression of bco1 or bco1l was induced by adding IPTG to a final concentration of 0.2 mM. After growing the cultures for an 16 h at 25 °C in darkness, bacteria were harvested by centrifugation. Carotenoid cleavage activity was visualized qualitatively by the lack of accumulated carotenoids which results in the absence of the yellow color characteristic of non-transformed control cells. Extraction of carotenoids and retinoids was conducted as previously described ^21^. All HPLC analyses were performed on a normal-phase Zorbax Sil (5 μm, 4.6 x 150 mm) column. Chromatographic separation was achieved by isocratic flow (1.4 ml/min) of a mixture of ethyl acetate and hexane. For the separation of the different carotenoids and retinoids, hexane was mixed with ethyl acetate in specific ratios for various time periods. For determination of carotenoids and retinoids, the HPLC column was scaled with known amounts of authentic standard substances.

### Homology modeling of salmon Bco1 and Bco1l

The multiple sequence alignment of Bco1 (and Atlantic salmon Bco1l) sequences from different species was carried out with the ClustalW program (http://www.ebi.ac.uk/clustalw/), and was manually modified afterwards. The three-dimensional structures of salmon Bco1 and Bco1l were generated through SWISS-PROT using RPE65 (PDB 3fsn.1.a) as the template, while PyMol (Schrödinger, LLC) was used for viewing.

## Author Contributions Statement

HH identified and characterised bco1 and bco1l homeologs in salmon. MS analysed the SNP-data (GWAS). NZ did the functional testing of the bco1/bco1l paralogs. MS, HH and NZ contributed to drafting the paper. JT supervised the functional testing together with JvL and DIV. FG did the RNAseq and analysed the data. JvL provided the E. Coli strain engineered to produce beta-caroten/zeaxanthin, and did the HPLC-studies. TM and SK contributed to the experimental design and provided samples and phenotype data for the GWAS. SL initiated and supervised the GWAS. All authors discussed and interpreted results, and contributed to writing of the paper. DIV made a final draft of the paper, and all authors read and approved the submitted manuscript.

## Acknowledgements

This project has received financial support from the Research Council of Norway, projects no. 177036 (Genofisk), 221734 (AquaGenome) and Norwegian University of Life Sciences (NMBU) and AquaGen ltd.

## Data Availability

The SNPs without a pre-existing rs-number has been submitted to European Variation Archive (EVA) under Project: PRJEB28258. All Illumina sequence reads are publicly accessible through the European Nucleotide Archive (http://www.ebi.ac.uk/ena) with the accession PRJEB10297. Additional datasets generated during and/or analysed during the current study are available from the corresponding author on reasonable request.

