## Supplemental information for "Genomic and functional gene studies suggest a key role of *beta-carotene oxygenase 1 like (bco1l)* gene in salmon flesh color"

### FIGURE LEGENDS

#### Figure S1

Protein sequence alignment of BCO1 from human, mice, rat, cattle, sheep and zebrafish together with Salmo salar Bco1 and Bco1l. The four histidines coordinating the $Fe^{2+}$ are shown in red letters. The additional residues making up the active site is highlighted with a black background, while the residues highlighted with a brown background are forming the patch located at the entrance to the active tunnel. The sequences are: NP_059125.2 (*Homo sapiens*), NP_001019730.1 (*Bos taurus*), NP_001295500.1 (*Ovis aries*), NP_067461.2 (*Mus musculus*), NP_446100.2 (*Rattus norvegicus*), NP_001315424.1 (*Danio rerio*), NP_001266000.1 (*Salmo salar - bco1*), NP_001266003.1 (*Salmo salar - bco1l*)

#### Figure S2

(A) Overlay cartoon view of predicted and spatially aligned Atlantic salmon Bco1 (purple) and Bco1l (cyan) structures, modeled with SWISS-MODEL program based on related RPE65 template 3fsn.1A (2.14 Å). (B) The iron cofactor, shown as an orange sphere, is directly coordinated by four histidine residues. (C) Iron cofactor with all the surrounding amino acid residues. For viewing PyMol was used.

#### Figure S3

Regions of misalignment between predicted 3D Bco1 and Bco1l models.

```
                            1         10         20         30         40
Homo sapiens_BCO1           ~MDIIFGRNR KEQLEPVRAK VTGKIPAWLQ GTLLRNGPGM HTVGESRYNH
Bos taurus_BCO1             ~MEIIFGRNK KEQLEPVRAR VTGKIPAWLQ GILLRNGPGM HTVGETRYNH
Ovis aries_BCO1             ~MEIIFGRNK KEQLEPVRAR VTGKIPAWLQ GTLLRNGPGM HTVGETRYNH
Mus musculus_BCO1           ~MEIIFGQNK KEQLEPVQAK VTGSIPAWLQ GTLLRNGPGM HTVGESKYNH
Rattus norvegicus_BCO1      ~MEIIFGRNK KEQLEPLRAT VTGSIPAWLQ GTLLRNGPGM HTVGDSKYNH
Danio rerio_BCO1            ~MQYDYGKNK EEHPEPIKTE VKGSIPEWVQ GTLIRNGPGM FSVGETTYNH
Salmo salar_BCO1            ~MAYDYAKNR EERPDPVKAD LKGNLPSWLQ GTLLRNGPGI FSVGDTTYNH
Salmo salar_BCO1l           MSQSLIGKNG TESPEPVKAE VTGCVPEWLQ GTLLRNGPGL FNVGATEYNH
                            50         60         70         80         90
Homo sapiens_BCO1           WFDGLALLHS FTIRDGEVYY RSKYLRSDTY NTNIEANRIV VSEFGTMAYP
Bos taurus_BCO1             WFDGLALLHS FTIRDGEVYY RSKYLRSDTY TANIEANRIV VSEFGTMAYP
Ovis aries_BCO1             WFDGLALLHS FTIRDGEVYY RSKYLRSDTY TANIEANRIV VSEFGTMAYP
Mus musculus_BCO1           WFDGLALLHS FSIRDGEVFY RSKYLQSDTY IANIEANRIV VSEFGTMAYP
Rattus norvegicus_BCO1      WFDGLALLHS FSIRDGEVFY RSKYLQSDTY NANIEANRIV VSEFGTMAYP
Danio rerio_BCO1            WFDGMALLHS FAINKGEVTY RSRYLRGDTY NSNMQANRIV VSEMGTMAYP
Salmo salar_BCO1            WFDGMALMHS FTFKDGEVIY RSRYLRGDTY KDNMAAKRIV VSEMGTMAYP
Salmo salar_BCO1l           WFDGMALIHS FTFKDGEVYY RSKFLRSDTF KKNTQANKIV VSEFGTMIYP
                            100        110        120        130        140
Homo sapiens_BCO1           DPCKNIFSKA FSYLSHTIPD FTDNCLINIM KCGEDFYATS ETNYIRKINP
Bos taurus_BCO1             DPCKNIFSKA FSYLSHTIPD FTDNCLINIR RCGEDFYATT ETSYIRRINP
Ovis aries_BCO1             DPCKNIFSKA FSYLSHTIPD FTDNCLINIM RCGEDFYATT ETNYIRKINP
Mus musculus_BCO1           DPCKNIFSKA FSYLSHTIPD FTDNCLINIM KCGEDFYATT ETNYIRKIDP
Rattus norvegicus_BCO1      DPCKNIFSKA FSYLSHTIPD FTDNCLINIM KCGEDFYATT ETNYIRKIDP
Danio rerio_BCO1            DPCKNIFSKV ITFLSHTIPD FTDNCGNNII KYGNDFHATS ETNYIRKIDP
Salmo salar_BCO1            DPGKNVISRV ITFLNHTVPD FTDNCGNNFI RYGKDYYATS ETNYIRKVDP
Salmo salar_BCO1l           DPCKNIFSKA FSYLLAAIPD FTDNNLINII RYGEDYYASS EVNYMNQIDP
                            150        160        170        180        190
Homo sapiens_BCO1           QTLETLEKVD YRKYVAVNLA TSHPHYDEAG NVLNMGTSIV EKGKTKYVIF
Bos taurus_BCO1             QTLETLEKVD FRKYVAVNLA TSHPHYDAAG NVLNVGTSIV DKGKTKYVIF
Ovis aries_BCO1             QTLETLEKVD YRKYVAVNLA TSHPHYDAAG NVLNVGTSIV DKGKTKYVIF
```

| | | | | | |
|---|---|---|---|---|---|
| *Mus musculus*_BCO1 | QTLETLEKVD | YRKYVAVNLA | TSHPHYDEAG | NVLNMGTSVV | DKGRTKYVIF |
| *Rattus norvegicus*_BCO1 | QTLETLEKVD | YRKYVAVNLA | TSHPHYDEAG | NVLNMGTSIA | DKGRTKYVMF |
| *Danio rerio*_BCO1 | VTLETQEKID | YLKYLPVSIV | ASHTHYDKEG | NSYSMGTCIA | EKGKTKYMLF |
| Salmo salar_BCO1 | VTLETQDKVD | YMKYLAVNLV | TSHPHYDKDG | TAYNIGTSIA | EKGKTKYTLF |
| Salmo salar_BCO1l | MTLDVIGKMN | YRNHIALNMA | TAHPHYDDEG | NTYNMGTALM | RFGMPNYVIF |
| | 200 | 210 | 220 | 230 | 240 |
| *Homo sapiens*_BCO1 | KIPATVPEGK | KQGKSPWKHT | EVFCSIPSRS | LLSPSYYHSF | GVTENYVIFL |
| *Bos taurus*_BCO1 | KIPAPVPGGR | KEGRSPLKDT | EVFCSIAAHS | LLSPSYYHSF | GVSENYIIFL |
| *Ovis aries*_BCO1 | KIPATVPGGR | KEGRSPLKDA | EVFCSIAARS | LLSPSYYHSF | GVTENYVVFL |
| *Mus musculus*_BCO1 | KIPATVPDSK | KKGKSPVKHA | EVFCSISSRS | LLSPSYYHSF | GVTENYVVFL |
| *Rattus norvegicus*_BCO1 | KIPATAPGSK | KKGKNPLKHS | EVFCSIPSRS | LLSPSYYHSF | GVTENYVVFL |
| *Danio rerio*_BCO1 | KVPG...ESR | PDGSPPLKSA | EAVCTLPCRS | LLTPSYYHSF | GMTDNYFIFI |
| Salmo salar_BCO1 | KVPDTTAGDK | ANASPALKNL | EVICTVPCRS | LLSPSYYHSF | GMTDNYLIFI |
| Salmo salar_BCO1l | KVPVNA.SDK | EHKKPALRKV | KQVCNIPIRS | TLFPSYFHSF | GMTENYIIFV |
| | 250 | 260 | 270 | 280 | 290 |
| *Homo sapiens*_BCO1 | EQPFRLDILK | MATAYIRRMS | WASCLAFHRE | EKTYIHIIDQ | RTRQPVQTKF |
| *Bos taurus*_BCO1 | EQPFKLDILK | MATAYIRGVS | WASCLAFHGE | DKTHIHIIDR | RTRKPVPTKY |
| *Ovis aries*_BCO1 | EQPFKLDILK | MATAYIRGVS | WASCLAFHGE | DKTHIHIIDR | RTRKPVLAKY |
| *Mus musculus*_BCO1 | EQPFKLDILK | MATAYMRGVS | WASCMSFDRE | DKTYIHIIDQ | RTRKPVPTKF |
| *Rattus norvegicus*_BCO1 | EQPFKLDILK | MATAYMRGVS | WASCMTFCKE | DKTYIHIIDQ | KTRKPVPTKF |
| *Danio rerio*_BCO1 | EQPLKLDILK | MATAYLRRVS | WASCMKFHPE | DSTLIHLIDR | NTKKEVATKF |
| Salmo salar_BCO1 | EQPFKLDILK | MATAYMRGVN | WASCLKFCPE | ENTLIHLIDR | KTGKEVGIKY |
| Salmo salar_BCO1l | EQPFKLDILR | LATAIFRRVT | WASCLKYDKE | DITLIHLIDK | KTGKAVSTKF |
| | 300 | 310 | 320 | 330 | 340 |
| *Homo sapiens*_BCO1 | YTDAMVVFHH | VNAYEEDGCI | VFDVIAYEDN | SLYQLFYLAN | LNQDFKE... |
| *Bos taurus*_BCO1 | HTDPMVVFHH | VNAYEEDGCL | LFDVITYEDG | SLYQLFYLAN | LNEDFKE... |
| *Ovis aries*_BCO1 | HTDPMVVFHH | VNAYEEDGCL | LFDVIAYEDG | SLYQLFYLAN | LNEDFKE... |
| *Mus musculus*_BCO1 | YTDPMVVFHH | VNAYEEDGCV | LFDVIAYEDS | SLYQLFYLAN | LNKDFEE... |
| *Rattus norvegicus*_BCO1 | YTDPMVVFHH | VNAYEEDGCV | LFDVIAYEDN | SLYQLFYLAN | LNKDFEE... |
| *Danio rerio*_BCO1 | YTDAMTVYHQ | VNAFEDDGHV | VFDVIAYDDN | NLYEFFYLNK | LKETMGA... |
| Salmo salar_BCO1 | YTEAMIVYHH | VNAFEEDGHV | IFDVIAYEDP | SLYNMFYLNV | LKEQSKA... |
| Salmo salar_BCO1l | YTDALVVFHH | INAYEDDGHV | VFDMITYKDG | NLYEMFYFAN | LRKETQEFIE |

| | | | | | |
|---|---|---|---|---|---|
| | | 356 | 366 | 376 | 386 |
| *Homo sapiens*_BCO1 | .NSRLTSVPT | LRRFAVPLHV | DKNAEVGTNL | IKVASTTATA | LKEEDGQVYC |
| *Bos taurus*_BCO1 | .NSRLTSMPT | LKRFVLPLHV | DKNAEVGSNL | IKLSSTTARA | LKEKDDQVYC |
| *Ovis aries*_BCO1 | .NSRLTSMPT | LKRFVLPLHV | DKNAEVGSNL | INLSSTTARA | LKEKDGQVYC |
| *Mus musculus*_BCO1 | .KSRLTSVPT | LRRFAVPLHV | DKDAEVGSNL | VKVSSTTATA | LKEKDGHVYC |
| *Rattus norvegicus*_BCO1 | .KSRLTSVPT | LRRFAVPLHV | DKDAEVGSNL | VKVSSTTATA | LKEKDDHVYC |
| *Danio rerio*_BCO1 | ..TNLYCKPK | FTRFVFPL.S | DQG.ETGENL | VKLKYTTASA | VKEKDGKIMC |
| Salmo salar_BCO1 | ...SAMSVPK | CKRFALPVQN | DKGIDVGDDM | VKLQYTTASA | VKEKEGKLLC |
| Salmo salar_BCO1l | SNKVNISPPI | CQRFVLPLTV | DKDTSNGTNL | VRLKDTTAKA | VMQSDGSLYC |
| | 396 | 406 | 416 | 426 | 436 |
| *Homo sapiens*_BCO1 | QPEFLYEGLE | LPRVNYAHNG | KQYRYVFATG | VQWSPIPTKI | IKYDILTKSS |
| *Bos taurus*_BCO1 | QPELLCEGLE | LPHINYAHNG | QPYRYIFAAG | VQWSPRPLIY | AAIR.LAKSS |
| *Ovis aries*_BCO1 | QPELLYEGLE | LPRINYAHNG | KPYRYVFAAG | VQWSPIPTQI | IKYDILTKSS |
| *Mus musculus*_BCO1 | QPEVLYEGLE | LPRINYAYNG | KPYRYIFAAE | VQWSPVPTKI | LKYDILTKSS |
| *Rattus norvegicus*_BCO1 | QPEVLYEGLE | LPRINYAHNG | KPYRYIFAAE | VQWSPVPTKI | LKYDVLTKSS |
| *Danio rerio*_BCO1 | QGEVLCEGVE | LPRINYNFNG | KKYRYSYMCC | VDESPVATRI | VKFDADTKQQ |
| Salmo salar_BCO1 | QPEVLCDGVE | LPRINYDFNG | KKYRFVYMTC | VAMSAVATKI | MKLDTETKER |
| Salmo salar_BCO1l | LPETIFQGLE | LPGMNYKFNG | KKYRYFYGSR | VEWTPHPNKI | GKVDIVTRKY |
| | 446 | 456 | 466 | 476 | 486 |
| *Homo sapiens*_BCO1 | LKWREDDCWP | AEPLFVPAPG | AKDEDDGVIL | SAIVSTDPQK | LPFLLILDAK |
| *Bos taurus*_BCO1 | LTWKEEHCWP | AEPLFVPTPG | AKDEDDGIIL | SAIVSTDPQK | SPFLLVLDAR |
| *Ovis aries*_BCO1 | LKWGEEHCWP | AEPLFVPTPG | AKDEDDGIIL | SAIVSTDPQK | SPFLLVLDAR |
| *Mus musculus*_BCO1 | LKWSEESCWP | AEPLFVPTPG | AKDEDDGVIL | SAIVSTDPQK | LPFLLILDAK |
| *Rattus norvegicus*_BCO1 | LKWSEESCWP | AEPLFVPTPG | AKDEDDGVIL | SAIISTDPQK | LPFLLILDAK |
| *Danio rerio*_BCO1 | IEWKGDDGFA | SEPVFIPRPG | AVDEDDGVVL | TVIINNKPLQ | GGFLLVLDAK |
| Salmo salar_BCO1 | TEWREENCWP | SEPVFIPRPN | GEGEDDGVVL | TTVINSNPGE | SGFILVLDAK |
| Salmo salar_BCO1l | IEWTEKDCYP | SEPVFVASPG | AVEEDDGVIL | TSVVSLNPKK | SPFMLVLNAK |
| | 496 | 506 | 516 | 526 | 536 |
| *Homo sapiens*_BCO1 | SFTELARASV | DVDMHMDLHG | LFITDMDWDT | KKQAASEEQR | DRASDCHGAP |
| *Bos taurus*_BCO1 | TFTELARASV | DVEMHLDFHG | LFIPDAGRDP | GKQAPSQEAP | ARAAAGRAAP |
| *Ovis aries*_BCO1 | TFTELARASI | DVEMHLDIHG | LFIPDAGWDL | GKQAPSREAP | ARAAAGRAAP |
| *Mus musculus*_BCO1 | SFTELARASV | DADMHLDLHG | LFIPDADWNA | VKQTPAETQE | VENSDHPTDP |

```
Rattus norvegicus_BCO1 SFTELARASV DVDMHLDLHG LFIPDAGWNA VKQTPAKTQE DENSDHPTGL
Danio rerio_BCO1       SFKEIARACL DVEIHMDMHG YFIPGSS~~~ ~~~~~~~~~~ ~~~~~~~~~~
Salmo salar_BCO1       SFKEVARAHV NAELHMDMHG YFIPMEN~~~ ~~~~~~~~~~ ~~~~~~~~~~
Salmo salar_BCO1l      TFEEIARASI DASIHLDLHG HFIPTQSTN~ ~~~~~~~~~~ ~~~~~~~~~~
                       546
Homo sapiens_BCO1      LT~~~~~~~~ ~~~~~~~~~~ ~~~~~~~~~~ ~~~~~~~~~~ ~~~~~~~~~~
Bos taurus_BCO1        RTDSLEALVL GTSSAQLTAV PAPGEGRESG PSFHFAHILS AAALSQNSET
Ovis aries_BCO1        QT~~~~~~~~ ~~~~~~~~~~ ~~~~~~~~~~ ~~~~~~~~~~ ~~~~~~~~~~
Mus musculus_BCO1      TA........ .......... PELSHSENDF TAGHGGSSL~ ~~~~~~~~~~
Rattus norvegicus_BCO1 TA........ .......... PGLGHGENDF TAGHGGKSL~ ~~~~~~~~~~
Danio rerio_BCO1       ~~~~~~~~~~ ~~~~~~~~~~ ~~~~~~~~~~ ~~~~~~~~~~ ~~~~~~~~~~
Salmo salar_BCO1       ~~~~~~~~~~ ~~~~~~~~~~ ~~~~~~~~~~ ~~~~~~~~~~ ~~~~~~~~~~
Salmo salar_BCO1l      ~~~~~~~~~~ ~~~~~~~~~~ ~~~~~~~~~~ ~~~~~~~~~~ ~~~~~~~~~~

Homo sapiens_BCO1      ~~
Bos taurus_BCO1        ET
Ovis aries_BCO1        ~~
Mus musculus_BCO1      ~~
Rattus norvegicus_BCO1 ~~
Danio rerio_BCO1       ~~
Salmo salar_BCO1       ~~
Salmo salar_BCO1l      ~~
```

Figur S1 - Helgeland et al.

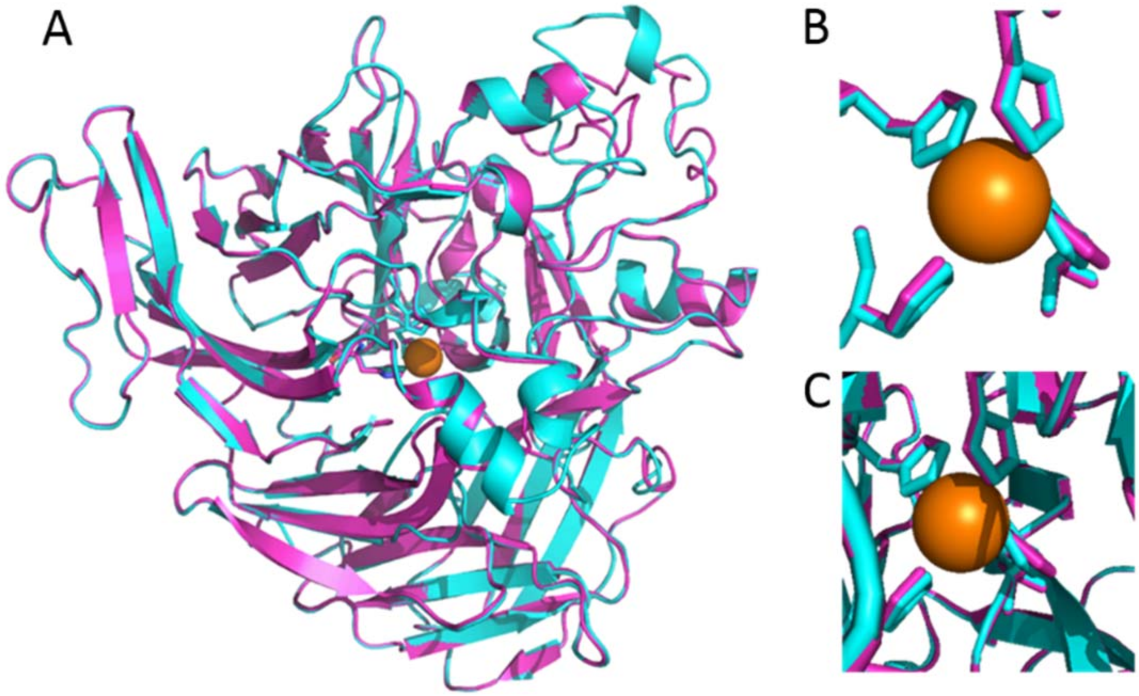


Figure S2 - Helgeland et al.

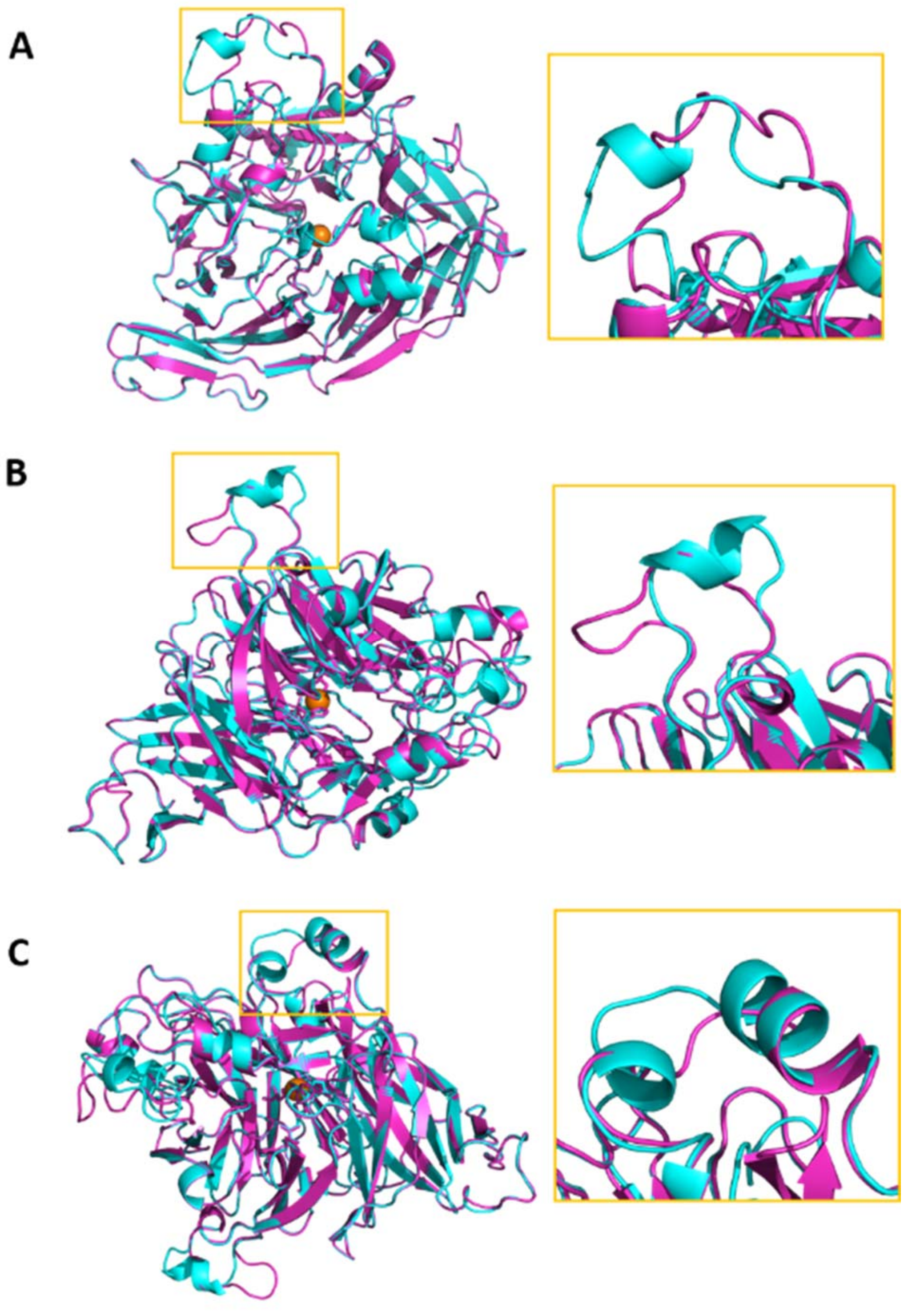


Figure S3 - Helgeland et al.

#### Table S1

SNP-identity, positions (assembly ICSASG_v2) and p-values for all the SNPs included in the fine-mapping at Ssa26. The SNPs without a rs-number has been submitted to European Variation Archive (EVA) under Project: PRJEB28258.

| SNP_ID | position | P-value | LogP |
|---|---|---|---|
| ctg7180001912552_1401_SCT | 18165496 | 1.330E-07 | 15.83 |
| rs863643757 | 18209026 | 1.563E-08 | 17.97 |
| rs863630607 | 18209503 | 4.376E-08 | 16.94 |
| rs864131335 | 18212528 | 3.846E-03 | 5.56 |
| rs863929219 | 18212572 | 1.229E-05 | 11.31 |
| rs863994564 | 18214352 | 7.186E-01 | 0.33 |
| rs863246859 | 18214880 | 4.850E-01 | 0.72 |
| rs863302789 | 18216152 | 5.767E-02 | 2.85 |
| rs864216165 | 18216446 | 5.725E-02 | 2.86 |
| rs864189383 | 18216795 | 5.767E-02 | 2.85 |
| rs864076812 | 18217233 | 9.143E-03 | 4.69 |
| rs863526953 | 18217539 | 5.725E-02 | 2.86 |
| rs863923796 | 18219658 | 5.767E-02 | 2.85 |
| rs863854450 | 18220244 | 5.767E-02 | 2.85 |
| rs863370817 | 18220852 | 6.305E-01 | 0.46 |
| rs863758464 | 18220868 | 6.305E-01 | 0.46 |
| rs863751717 | 18225298 | 5.192E-02 | 2.96 |
| rs863472796 | 18226558 | 5.811E-01 | 0.54 |
| rs863870255 | 18245647 | 1.128E-02 | 4.48 |
| rs863670640 | 18249825 | 3.691E-01 | 1.00 |
| rs863942373 | 18250502 | 7.280E-01 | 0.32 |
| rs863848334 | 18250708 | 5.951E-01 | 0.52 |
| rs863358817 | 18252043 | 5.246E-11 | 23.67 |
| rs863749263 | 18254032 | 7.606E-05 | 9.48 |
| rs863527645 | 18254565 | 6.628E-02 | 2.71 |
| rs863742312 | 18254876 | 6.628E-02 | 2.71 |
| rs863875634 | 18255060 | 3.691E-01 | 1.00 |
| rs864212792 | 18255144 | 2.818E-04 | 8.17 |
| rs863445911 | 18255574 | 2.097E-04 | 8.47 |
| rs863279327 | 18255833 | 3.450E-04 | 7.97 |
| rs863668595 | 18255912 | 1.100E-11 | 25.23 |
| rs863405534 | 18256181 | 4.237E-04 | 7.77 |
| rs863678353 | 18257349 | 9.904E-13 | 27.64 |
| rs864011705 | 18257454 | 9.904E-13 | 27.64 |

| | | | |
|---|---|---|---|
| rs863268852 | 18259051 | 6.802E-02 | 2.69 |
| rs864146797 | 18278051 | 6.598E-01 | 0.42 |
| rs864022302 | 18281310 | 1.424E-02 | 4.25 |
| rs863861608 | 18285252 | 1.222E-06 | 13.61 |
| rs863645922 | 18285350 | 3.452E-07 | 14.88 |
| rs864159650 | 18286456 | 3.244E-01 | 1.13 |
| rs864062338 | 18286948 | 1.084E-02 | 4.52 |
| rs863545505 | 18287102 | 8.816E-01 | 0.13 |
| rs863742875 | 18287250 | 8.816E-01 | 0.13 |
| rs864214566 | 18287506 | 3.691E-01 | 1.00 |
| rs863530134 | 18287701 | 3.691E-01 | 1.00 |
| rs863610917 | 18289359 | 3.421E-06 | 12.59 |
| rs863809717 | 18297483 | 1.215E-03 | 6.71 |
| ctg7180001937482_1652_SAG | 18300048 | 9.065E-01 | 0.10 |
| rs863570483 | 18330028 | 6.889E-01 | 0.37 |
| rs863460169 | 18332201 | 6.807E-04 | 7.29 |
| rs864222221 | 18332544 | 1.388E-02 | 4.28 |
| rs863239536 | 18339421 | 2.704E-01 | 1.31 |
| rs863261398 | 18344217 | 2.225E-03 | 6.11 |
| rs864045796 | 18344416 | 2.225E-03 | 6.11 |
| rs863467318 | 18347696 | 2.225E-03 | 6.11 |
| rs863848265 | 18352177 | 3.486E-01 | 1.05 |
| rs863951906 | 18357758 | 1.298E-02 | 4.34 |
| rs863883327 | 18366158 | 1.138E-02 | 4.48 |
| rs863409857 | 18367282 | 9.249E-02 | 2.38 |
| rs863756163 | 18369434 | 9.249E-02 | 2.38 |
| rs863621526 | 18372945 | 1.173E-01 | 2.14 |
| rs863257497 | 18373077 | 1.173E-01 | 2.14 |
| rs863930762 | 18374240 | 1.173E-01 | 2.14 |
| rs863955152 | 18374323 | 5.533E-02 | 2.89 |
| rs863270876 | 18375356 | 1.703E-01 | 1.77 |
| rs864226610 | 18375430 | 6.788E-02 | 2.69 |
| rs863405630 | 18376136 | 1.704E-01 | 1.77 |
| rs863490287 | 18376340 | 2.978E-02 | 3.51 |
| rs864231524 | 18376749 | 2.978E-02 | 3.51 |
| rs863551430 | 18380372 | 3.378E-04 | 7.99 |
| rs863245462 | 18383378 | 9.139E-02 | 2.39 |
| rs863242518 | 18383929 | 5.039E-01 | 0.69 |
| rs863504373 | 18385547 | 6.855E-01 | 0.38 |
| rs864004026 | 18396882 | 5.578E-01 | 0.58 |
| ctg7180001910618_6988_SAG | 18397478 | 7.997E-01 | 0.22 |
| rs863904970 | 18400408 | 3.476E-01 | 1.06 |
| rs863883651 | 18400771 | 2.341E-05 | 10.66 |

| | | | |
|---|---|---|---|
| rs863655491 | 18401061 | 3.476E-01 | 1.06 |
| rs863336140 | 18402164 | 8.303E-06 | 11.70 |
| rs863827406 | 18407360 | 1.844E-01 | 1.69 |
| ctg7180001934204_510_SAG | 18411761 | 4.158E-03 | 5.48 |
| rs864211565 | 18412415 | 6.664E-02 | 2.71 |
| ctg7180001934204_1641_SAG | 18412892 | 3.667E-05 | 10.21 |
| rs863739786 | 18414556 | 3.844E-03 | 5.56 |
| rs159401219 | 18414602 | 1.621E-06 | 13.33 |
| rs159401218 | 18414949 | 4.128E-02 | 3.19 |
| rs863879353 | 18415545 | 1.091E-02 | 4.52 |
| rs864113451 | 18415688 | 9.424E-10 | 20.78 |
| rs863601299 | 18415730 | 3.001E-01 | 1.20 |
| rs863898337 | 18415786 | 5.117E-13 | 28.30 |
| rs863905018 | 18422579 | 3.542E-01 | 1.04 |
| rs864138807 | 18423344 | 5.101E-04 | 7.58 |
| rs864120442 | 18424569 | 9.743E-04 | 6.93 |
| rs863373926 | 18426927 | 1.318E-02 | 4.33 |
| rs863348554 | 18427812 | 2.370E-02 | 3.74 |
| ctg7180001866800_6544_SAG | 18429158 | 7.757E-01 | 0.25 |
| rs863402331 | 18432173 | 7.470E-08 | 16.41 |
| rs864029623 | 18432240 | 3.006E-01 | 1.20 |
| rs863661748 | 18432582 | 1.693E-07 | 15.59 |
| rs863711429 | 18432785 | 6.071E-09 | 18.92 |
| rs863291822 | 18432840 | 1.527E-03 | 6.48 |
| rs863564784 | 18432868 | 3.015E-01 | 1.20 |
| rs863267812 | 18433531 | 3.015E-01 | 1.20 |
| rs863746472 | 18433823 | 1.331E-04 | 8.92 |
| rs863419768 | 18433908 | 1.691E-04 | 8.69 |
| rs863235361 | 18433914 | 1.691E-04 | 8.69 |
| rs863315971 | 18434126 | 1.029E-07 | 16.09 |
| rs863274339 | 18434589 | 1.691E-04 | 8.69 |
| ctg7180001934204_23777_SAG | 18435029 | 5.895E-02 | 2.83 |
| rs864091273 | 18435996 | 4.530E-09 | 19.21 |
| rs863689986 | 18436349 | 4.820E-02 | 3.03 |
| rs863933324 | 18436862 | 7.296E-01 | 0.32 |
| rs864043929 | 18440020 | 2.569E-01 | 1.36 |
| rs863746292 | 18440590 | 2.569E-01 | 1.36 |
| rs864000441 | 18441078 | 2.569E-01 | 1.36 |
| rs863630746 | 18441401 | 3.414E-02 | 3.38 |
| rs863740261 | 18443283 | 2.569E-01 | 1.36 |
| rs863974888 | 18447307 | 1.341E-04 | 8.92 |
| rs863470605 | 18447523 | 2.569E-01 | 1.36 |
| rs864216526 | 18449231 | 5.939E-01 | 0.52 |

| | | | |
|---|---|---|---|
| ctg7180001841303_5259_SAG | 18457743 | 1.005E-02 | 4.60 |
| rs863337839 | 18459452 | 2.824E-01 | 1.26 |
| rs864145384 | 18460914 | 1.389E-06 | 13.49 |
| rs863967569 | 18461912 | 6.130E-02 | 2.79 |
| ctg7180001352914_8470_SGT | 18467059 | 9.203E-07 | 13.90 |
| rs863996077 | 18467277 | 1.128E-07 | 16.00 |
| ctg7180001352914_9263_SAG | 18467852 | 8.782E-05 | 9.34 |
| rs863921815 | 18479105 | 3.013E-01 | 1.20 |
| rs863707711 | 18479426 | 3.013E-01 | 1.20 |
| rs863536299 | 18485339 | 3.029E-03 | 5.80 |
| rs863473627 | 18487418 | 8.131E-11 | 23.23 |
| rs863324020 | 18487479 | 5.521E-02 | 2.90 |
| rs863521503 | 18488034 | 7.304E-02 | 2.62 |
| rs864141582 | 18490224 | 4.953E-04 | 7.61 |
| rs863934226 | 18492896 | 6.967E-06 | 11.87 |
| rs863584931 | 18504183 | 2.969E-02 | 3.52 |
| rs863768234 | 18504237 | 1.259E-04 | 8.98 |
| rs863854470 | 18504285 | 4.283E-02 | 3.15 |
| rs864081920 | 18506731 | 2.218E-04 | 8.41 |
| rs863704000 | 18508611 | 2.218E-04 | 8.41 |
| rs864075902 | 18514528 | 2.218E-04 | 8.41 |
| rs863585735 | 18514538 | 2.218E-04 | 8.41 |
| rs863454153 | 18518899 | 1.259E-04 | 8.98 |
| rs863606098 | 18522330 | 2.909E-04 | 8.14 |
| rs863247952 | 18522988 | 2.836E-02 | 3.56 |
| rs863518137 | 18523082 | 9.840E-01 | 0.02 |
| rs863670279 | 18524388 | 3.036E-02 | 3.49 |
| rs864180255 | 18525089 | 6.465E-01 | 0.44 |
| rs863525606 | 18525582 | 4.475E-04 | 7.71 |
| rs863597426 | 18530653 | 6.698E-02 | 2.70 |
| rs864066718 | 18531084 | 8.904E-01 | 0.12 |
| rs863232239 | 18532566 | 8.904E-01 | 0.12 |
| rs863633278 | 18533037 | 8.969E-03 | 4.71 |
| rs863848436 | 18533819 | 8.232E-06 | 11.71 |
| rs863786003 | 18534062 | 7.277E-01 | 0.32 |
| ctg7180001805095_6306_SAC | 18542907 | 1.463E-01 | 1.92 |
| rs863695107 | 18547947 | 5.617E-02 | 2.88 |
| rs863325783 | 18548016 | 6.456E-01 | 0.44 |
| rs863967251 | 18549631 | 4.535E-02 | 3.09 |
| rs863870456 | 18550971 | 5.803E-05 | 9.75 |
| rs864052330 | 18551004 | 5.803E-05 | 9.75 |
| rs863879095 | 18554315 | 7.325E-01 | 0.31 |
| rs863866399 | 18555608 | 7.325E-01 | 0.31 |

| | | | |
|---|---|---|---|
| rs863603382 | 18556374 | 6.253E-01 | 0.47 |
| rs863521067 | 18569929 | 7.655E-02 | 2.57 |
| rs864180301 | 18573106 | 1.937E-05 | 10.85 |
| rs863428566 | 18579728 | 4.974E-07 | 14.51 |
| rs863629571 | 18579732 | 6.780E-05 | 9.60 |
| rs863512936 | 18581586 | 1.502E-03 | 6.50 |
| rs864142918 | 18583319 | 2.591E-03 | 5.96 |
| rs863432906 | 18583940 | 6.471E-01 | 0.44 |
| rs863823788 | 18584663 | 6.471E-01 | 0.44 |
| rs863565691 | 18586336 | 3.828E-05 | 10.17 |
| rs863562206 | 18586597 | 4.954E-01 | 0.70 |
| rs863815715 | 18599967 | 2.483E-06 | 12.91 |
| rs863749953 | 18600666 | 6.667E-01 | 0.41 |
| rs863483118 | 18600892 | 2.659E-08 | 17.44 |
| rs863294192 | 18601824 | 2.659E-08 | 17.44 |
| rs863372498 | 18607960 | 5.428E-02 | 2.91 |
| rs864247432 | 18608983 | 2.659E-08 | 17.44 |
| rs863470433 | 18609040 | 2.659E-08 | 17.44 |
| rs863799208 | 18609181 | 2.659E-08 | 17.44 |
| rs863354021 | 18619054 | 2.154E-02 | 3.84 |
| rs864228317 | 18622171 | 4.925E-01 | 0.71 |
| rs864176083 | 18622336 | 2.209E-01 | 1.51 |
| rs864083818 | 18622816 | 6.213E-04 | 7.38 |
| rs863257025 | 18622998 | 4.552E-06 | 12.30 |
| rs863276828 | 18624465 | 1.828E-02 | 4.00 |
| rs864022708 | 18624688 | 2.452E-02 | 3.71 |
| ctg7180001779364_1565_SCT | 18624729 | 1.231E-02 | 4.40 |
| ctg7180001860735_7383_SAG | 18624863 | 2.138E-02 | 3.85 |
| rs863438371 | 18625071 | 2.152E-07 | 15.35 |
| rs863734473 | 18625261 | 3.586E-05 | 10.24 |
| rs864057710 | 18625584 | 2.467E-03 | 6.00 |
| rs863659014 | 18627218 | 1.150E-06 | 13.68 |
| rs863737351 | 18627328 | 3.586E-05 | 10.24 |
| rs864074323 | 18627815 | 3.754E-01 | 0.98 |
| rs864063606 | 18628106 | 1.744E-05 | 10.96 |
| rs863607845 | 18629160 | 3.754E-01 | 0.98 |
| rs863365385 | 18629283 | 3.680E-07 | 14.82 |
| rs864109729 | 18634188 | 6.154E-06 | 12.00 |
| ctg7180001334602_552_SCT | 18635466 | 5.000E-01 | 0.69 |
| rs863457882 | 18637926 | 6.154E-06 | 12.00 |
| rs864036713 | 18638069 | 2.659E-08 | 17.44 |
| rs863427677 | 18638690 | 2.659E-08 | 17.44 |
| rs863498848 | 18638885 | 1.439E-01 | 1.94 |

| | | | |
|---|---|---|---|
| rs863293399 | 18643466 | 6.714E-01 | 0.40 |
| rs863664652 | 18643877 | 7.235E-09 | 18.74 |
| rs863900687 | 18644168 | 3.080E-06 | 12.69 |
| rs864243805 | 18644202 | 2.659E-08 | 17.44 |
| rs864105311 | 18644640 | 2.659E-08 | 17.44 |
| rs863740510 | 18648223 | 2.659E-08 | 17.44 |
| rs863308265 | 18648722 | 4.132E-03 | 5.49 |
| rs863795279 | 18648849 | 2.659E-08 | 17.44 |
| rs863422146 | 18649630 | 2.659E-08 | 17.44 |
| rs863956787 | 18649744 | 2.659E-08 | 17.44 |
| rs863538184 | 18649792 | 1.127E-01 | 2.18 |
| rs864031980 | 18650133 | 1.341E-07 | 15.82 |
| rs863418676 | 18670222 | 1.270E-04 | 8.97 |
| rs863252336 | 18670508 | 6.560E-01 | 0.42 |
| rs863883261 | 18671558 | 1.879E-01 | 1.67 |
| rs864119264 | 18672365 | 1.879E-01 | 1.67 |
| rs863346645 | 18672938 | 1.154E-07 | 15.97 |
| ctg7180001901066_4965_SAT | 18673463 | 8.571E-06 | 11.67 |
| rs864224668 | 18675116 | 2.307E-04 | 8.37 |
| rs863946442 | 18675437 | 2.362E-05 | 10.65 |
| rs863373340 | 18675525 | 2.860E-04 | 8.16 |
| rs864230082 | 18675920 | 2.659E-08 | 17.44 |
| rs864087201 | 18676038 | 2.659E-08 | 17.44 |
| rs863787462 | 18676433 | 7.306E-03 | 4.92 |
| rs863838924 | 18676485 | 6.563E-05 | 9.63 |
| rs863639625 | 18677056 | 1.816E-04 | 8.61 |
| rs863734750 | 18677555 | 1.909E-05 | 10.87 |
| rs863264291 | 18678887 | 1.290E-07 | 15.86 |
| rs863710116 | 18678948 | 1.356E-02 | 4.30 |
| rs863850827 | 18679128 | 1.356E-02 | 4.30 |
| rs863401956 | 18690950 | 2.495E-06 | 12.90 |
| rs863860980 | 18694935 | 5.355E-02 | 2.93 |
| rs864229410 | 18695400 | 6.317E-01 | 0.46 |
| rs864097358 | 18710902 | 2.000E-08 | 17.73 |
| rs863568638 | 18710943 | 1.869E-05 | 10.89 |
| rs863772437 | 18711924 | 8.813E-01 | 0.13 |
| rs863408412 | 18712894 | 8.813E-01 | 0.13 |
| rs864191959 | 18713145 | 8.813E-01 | 0.13 |
| rs863747597 | 18715000 | 8.813E-01 | 0.13 |
| rs863697357 | 18715771 | 2.698E-03 | 5.92 |
| rs863775212 | 18716559 | 2.659E-08 | 17.44 |
| rs864187056 | 18716576 | 2.659E-08 | 17.44 |
| rs863525292 | 18717095 | 5.745E-01 | 0.55 |

| | | | |
|---|---|---|---|
| rs864064286 | 18717423 | 3.412E-06 | 12.59 |
| rs863661189 | 18717608 | 3.996E-05 | 10.13 |
| ctg7180001895681_5855_SAC | 18718235 | 9.121E-01 | 0.09 |
| rs863874948 | 18718773 | 5.959E-08 | 16.64 |
| rs864156042 | 18720538 | 5.110E-01 | 0.67 |
| rs864226477 | 18720593 | 5.110E-01 | 0.67 |
| rs863860690 | 18725688 | 5.469E-02 | 2.91 |
| rs863333509 | 18729684 | 9.274E-01 | 0.08 |
| rs863850490 | 18733202 | 2.659E-08 | 17.44 |
| rs864152410 | 18737057 | 2.659E-08 | 17.44 |
| rs863241295 | 18738280 | 2.443E-03 | 6.01 |
| rs864207151 | 18738817 | 2.443E-03 | 6.01 |
| rs863664674 | 18738880 | 1.799E-08 | 17.83 |
| rs864225571 | 18739096 | 1.412E-01 | 1.96 |
| rs864124247 | 18739626 | 6.994E-01 | 0.36 |
| rs863983453 | 18739795 | 3.169E-04 | 8.06 |
| rs863243102 | 18740287 | 3.767E-01 | 0.98 |
| rs863781128 | 18740394 | 4.560E-04 | 7.69 |
| ctg7180001340274_15139_SGT | 18740601 | 9.609E-01 | 0.04 |
| rs863788280 | 18741600 | 9.601E-01 | 0.04 |
| rs864222418 | 18742780 | 1.908E-07 | 15.47 |
| rs863406367 | 18742930 | 8.948E-01 | 0.11 |
| rs863652269 | 18742958 | 1.386E-07 | 15.79 |
| rs863737359 | 18743190 | 1.725E-04 | 8.67 |
| rs864185737 | 18743261 | 1.725E-04 | 8.67 |
| rs863704627 | 18743932 | 1.412E-04 | 8.87 |
| rs863534823 | 18744097 | 1.412E-04 | 8.87 |
| rs863302884 | 18744580 | 8.516E-01 | 0.16 |
| rs863750215 | 18745576 | 1.412E-04 | 8.87 |
| rs864003384 | 18746919 | 1.412E-04 | 8.87 |
| rs863555760 | 18752404 | 5.161E-05 | 9.87 |
| rs863293163 | 18755881 | 4.227E-02 | 3.16 |
| rs863691131 | 18756307 | 5.320E-03 | 5.24 |
| rs864181485 | 18757599 | 7.067E-04 | 7.25 |
| rs864214258 | 18763582 | 1.762E-01 | 1.74 |
| rs864246016 | 18764397 | 2.708E-04 | 8.21 |
| rs863552849 | 18764947 | 3.336E-03 | 5.70 |
| rs863955128 | 18769453 | 9.221E-03 | 4.69 |
| ctg7180001328514_773_SGT | 18782635 | 4.834E-02 | 3.03 |
| rs864252727 | 18782806 | 9.822E-01 | 0.02 |
| rs863360388 | 18785547 | 9.822E-01 | 0.02 |
| rs863548835 | 18787338 | 1.005E-02 | 4.60 |
| rs863407086 | 18788399 | 1.005E-02 | 4.60 |

| | | | |
|---|---|---|---|
| rs864209010 | 18788474 | 2.122E-01 | 1.55 |
| rs863531362 | 18789091 | 1.005E-02 | 4.60 |
| rs863387654 | 18789510 | 1.005E-02 | 4.60 |
| rs863986381 | 18794987 | 1.461E-01 | 1.92 |
| rs863739822 | 18801802 | 1.005E-02 | 4.60 |
| ctg7180001744125_623_SCT | 18823705 | 1.200E-02 | 4.42 |
| rs864138462 | 18827172 | 2.724E-01 | 1.30 |
| rs864148002 | 18828430 | 7.029E-01 | 0.35 |
| rs863448305 | 18837393 | 6.100E-01 | 0.49 |
| rs864170707 | 18837668 | 6.100E-01 | 0.49 |
| rs863918410 | 18837868 | 7.088E-01 | 0.34 |
| rs863486915 | 18838454 | 5.416E-01 | 0.61 |
| rs863392682 | 18841031 | 4.051E-01 | 0.90 |
| rs863633225 | 18845870 | 8.576E-04 | 7.06 |
| rs863861104 | 18846924 | 8.576E-04 | 7.06 |
| rs863508427 | 18850773 | 1.343E-05 | 11.22 |
| rs863919979 | 18850805 | 1.343E-05 | 11.22 |
| rs863541055 | 18851187 | 2.067E-01 | 1.58 |
| rs863819255 | 18864180 | 5.702E-01 | 0.56 |
| rs863703516 | 18865539 | 1.184E-01 | 2.13 |
| rs863985900 | 18866647 | 1.184E-01 | 2.13 |
| rs863284245 | 18866942 | 1.184E-01 | 2.13 |
| rs863242467 | 18872043 | 1.184E-01 | 2.13 |
| ctg7180001805095_19769_SCT | 18881719 | 3.959E-03 | 5.53 |
| rs863836278 | 18882170 | 3.691E-01 | 1.00 |
| rs863981730 | 18888084 | 1.184E-01 | 2.13 |
| rs863918510 | 18888554 | 8.522E-01 | 0.16 |
| rs864029155 | 18889161 | 1.184E-01 | 2.13 |
| rs863229108 | 18889269 | 1.184E-01 | 2.13 |
| rs863320925 | 18890667 | 4.504E-05 | 10.01 |
| rs864098139 | 18891332 | 8.707E-01 | 0.14 |
| rs863291655 | 18891601 | 8.707E-01 | 0.14 |
| ctg7180001805095_14378_SCT | 18891696 | 3.988E-01 | 0.92 |
| rs863792522 | 18891905 | 8.707E-01 | 0.14 |
| rs864205400 | 18894521 | 4.409E-03 | 5.42 |
| rs863317802 | 18896131 | 4.275E-02 | 3.15 |
| rs864118606 | 18900481 | 2.965E-02 | 3.52 |
| rs863942301 | 18905202 | 2.965E-02 | 3.52 |
| rs863458548 | 18905614 | 2.965E-02 | 3.52 |
| rs863276889 | 18905782 | 2.965E-02 | 3.52 |
| rs863735591 | 18906569 | 5.706E-01 | 0.56 |
| rs864251506 | 18907256 | 8.545E-12 | 25.49 |
| rs863900233 | 18913428 | 2.079E-03 | 6.18 |

| | | | |
|---|---|---|---|
| rs863350932 | 18916170 | 2.079E-03 | 6.18 |
| rs863279457 | 18926749 | 3.387E-02 | 3.39 |
| rs863911761 | 18928361 | 3.127E-01 | 1.16 |
| rs864232964 | 18944506 | 1.222E-01 | 2.10 |
| rs864021810 | 18945934 | 1.222E-01 | 2.10 |
| rs863609169 | 18950641 | 1.080E-01 | 2.23 |
| rs864111848 | 18950916 | 7.442E-09 | 18.72 |
| rs863873516 | 18952527 | 7.442E-09 | 18.72 |
| rs863962390 | 18952568 | 3.860E-02 | 3.25 |
| rs863245319 | 18970280 | 3.241E-01 | 1.13 |
| ctg7180001467069_2463_SAC | 18972289 | 3.508E-01 | 1.05 |
| rs863962816 | 18987598 | 3.230E-06 | 12.64 |
| rs864173761 | 18987665 | 3.230E-06 | 12.64 |
| rs864111198 | 18987814 | 8.533E-02 | 2.46 |
| rs863244357 | 18988111 | 8.533E-02 | 2.46 |
| rs863342759 | 18989829 | 1.618E-04 | 8.73 |
| rs159406129 | 18990193 | 1.798E-02 | 4.02 |
| rs159406129 | 18990193 | 1.404E-02 | 4.27 |
| ctg7180001779364_3569_SAG | 18990295 | 1.231E-02 | 4.40 |
| rs863725961 | 18990715 | 4.842E-02 | 3.03 |
| rs863579211 | 18992195 | 3.531E-02 | 3.34 |
| rs864108707 | 18993133 | 1.231E-02 | 4.40 |
| rs864083648 | 18997024 | 1.708E-04 | 8.68 |
| rs864029813 | 18997358 | 3.531E-02 | 3.34 |
| rs864104224 | 18997416 | 2.227E-01 | 1.50 |
| rs863603320 | 19001155 | 1.405E-04 | 8.87 |
| rs864222936 | 19001953 | 1.405E-04 | 8.87 |
| rs863798621 | 19001990 | 2.134E-06 | 13.06 |
| rs863455321 | 19002100 | 1.405E-04 | 8.87 |
| rs863305376 | 19002245 | 1.405E-04 | 8.87 |
| rs864170292 | 19002795 | 2.155E-06 | 13.05 |
| rs863313903 | 19002968 | 3.531E-02 | 3.34 |
| rs863278458 | 19003453 | 3.531E-02 | 3.34 |
| rs863756893 | 19005565 | 2.141E-04 | 8.45 |
| rs863415577 | 19006593 | 7.171E-07 | 14.15 |
| rs863396798 | 19007006 | 2.157E-05 | 10.74 |
| rs864135251 | 19007154 | 7.171E-07 | 14.15 |
| rs864113346 | 19007446 | 1.980E-06 | 13.13 |
| rs864208778 | 19007511 | 3.531E-02 | 3.34 |
| rs864208778 | 19007511 | 3.531E-02 | 3.34 |
| BCMO1_81837 | 19008130 | 1.708E-04 | 8.68 |
| BCMO1_81820 | 19008147 | 3.531E-02 | 3.34 |
| rs863908213 | 19008790 | 2.917E-06 | 12.74 |

| | | | |
|---|---|---|---|
| rs863289778 | 19009217 | 2.119E-06 | 13.06 |
| rs863613538 | 19009726 | 1.632E-03 | 6.42 |
| rs863831492 | 19013109 | 4.694E-06 | 12.27 |
| BCMO1_SSA26MOD_422424 | 19017529 | 4.036E-01 | 0.91 |
| rs863772928 | 19018616 | 3.631E-02 | 3.32 |
| BCMO1_SSA26MOD_416513 | 19023616 | 4.267E-02 | 3.15 |
| rs863673647 | 19024187 | 2.024E-04 | 8.51 |
| rs863673647 | 19024187 | 7.289E-06 | 11.83 |
| rs863723023 | 19029542 | 7.289E-06 | 11.83 |
| rs864246870 | 19029622 | 7.289E-06 | 11.83 |
| rs863699183 | 19032982 | 4.317E-03 | 5.45 |
| rs863429657 | 19033176 | 4.317E-03 | 5.45 |
| BCMO1_ssa26mod_408753 | 19033212 | 3.132E-02 | 3.46 |
| rs863650429 | 19033344 | 6.409E-06 | 11.96 |
| BCMO1_SSA26MOD_407073 | 19034892 | 2.606E-04 | 8.25 |
| rs863841221 | 19035282 | 1.677E-11 | 24.81 |
| rs863735013 | 19039889 | 7.183E-04 | 7.24 |
| rs864185097 | 19042770 | 1.578E-04 | 8.75 |
| rs864185097 | 19042770 | 7.535E-06 | 11.80 |
| rs863815058 | 19043542 | 1.578E-04 | 8.75 |
| rs863815058 | 19043542 | 7.535E-06 | 11.80 |
| BCMO1_45633 | 19044104 | 2.418E-02 | 3.72 |
| rs863457647 | 19044305 | 1.578E-04 | 8.75 |
| ctg7180001733669_1558_SCT | 19047746 | 3.863E-02 | 3.25 |
| BCMO1_43314 | 19048254 | 1.530E-01 | 1.88 |
| rs864136920 | 19049331 | 1.434E-25 | 57.20 |
| rs863699640 | 19050133 | 1.094E-24 | 55.17 |
| rs159403238 | 19050343 | 1.434E-25 | 57.20 |
| rs159403238 | 19050343 | 1.434E-25 | 57.20 |
| BCMO1Jan2012_50882 | 19051049 | 4.762E-01 | 0.74 |
| rs863433469 | 19051075 | 1.530E-01 | 1.88 |
| BCMO1Jan2012_50796 | 19051135 | 1.179E-01 | 2.14 |
| rs864240256 | 19051152 | 1.201E-01 | 2.12 |
| ctg7180001733669_5231_SAG | 19051419 | 1.530E-01 | 1.88 |
| rs863294679 | 19051426 | 1.530E-01 | 1.88 |
| rs863460708 | 19052008 | 1.434E-25 | 57.20 |
| rs863979260 | 19052145 | 1.530E-01 | 1.88 |
| rs864121002 | 19053125 | 6.565E-01 | 0.42 |
| rs863622198 | 19053298 | 1.179E-01 | 2.14 |
| rs864133094 | 19053468 | 1.094E-24 | 55.17 |
| BCMO1_37970 | 19053598 | 1.179E-01 | 2.14 |
| rs864140717 | 19056436 | 1.179E-01 | 2.14 |
| rs863667798 | 19058667 | 1.530E-01 | 1.88 |

| | | | |
|---|---|---|---|
| rs863281847 | 19060211 | 3.290E-26 | 58.68 |
| rs863418153 | 19061809 | 1.719E-23 | 52.42 |
| rs863556042 | 19062953 | 3.574E-27 | 60.90 |
| rs863721927 | 19063311 | 1.188E-01 | 2.13 |
| rs863434642 | 19064216 | 2.946E-26 | 58.79 |
| rs863645576 | 19064952 | 5.023E-27 | 60.56 |
| rs863699886 | 19068122 | 6.677E-27 | 60.27 |
| BCMO1_SSA26MOD_361489 | 19068133 | 6.677E-27 | 60.27 |
| rs864032545 | 19069882 | 1.887E-27 | 61.53 |
| rs863388734 | 19069884 | 1.426E-28 | 64.12 |
| rs863560551 | 19073497 | 9.160E-30 | 66.86 |
| rs863946832 | 19076391 | 7.168E-01 | 0.33 |
| rs863532363 | 19077089 | 4.815E-27 | 60.60 |
| rs863554585 | 19077273 | 1.107E-34 | 78.19 |
| rs863442990 | 19079127 | 8.095E-54 | 122.25 |
| rs863867200 | 19079917 | 6.768E-35 | 78.68 |
| rs864103853 | 19080027 | 1.196E-09 | 20.54 |
| rs863785818 | 19081573 | 1.377E-53 | 121.72 |
| rs863641555 | 19082257 | 1.503E-11 | 24.92 |
| rs863758878 | 19082481 | 1.503E-11 | 24.92 |
| BCMO1_3801 | 19082663 | 1.153E-29 | 66.63 |
| BCMO1_3484 | 19082980 | 8.757E-24 | 53.09 |
| rs863682291 | 19083074 | 3.877E-11 | 23.97 |
| BCMO1_3241 | 19083223 | 2.259E-02 | 3.79 |
| rs863940219 | 19083998 | 9.841E-03 | 4.62 |
| BCMO1_1017 | 19085447 | 3.895E-01 | 0.94 |
| rs863480295 | 19085953 | 3.895E-01 | 0.94 |
| rs864104703 | 19086549 | 3.920E-01 | 0.94 |
| rs863997987 | 19086803 | 6.487E-01 | 0.43 |
| rs864163658 | 19088480 | 3.540E-16 | 35.58 |
| rs864239694 | 19090689 | 2.229E-01 | 1.50 |
| rs863356149 | 19092929 | 3.540E-16 | 35.58 |
| rs863242077 | 19093101 | 3.540E-16 | 35.58 |
| rs863837258 | 19093286 | 3.368E-01 | 1.09 |
| rs863560128 | 19093347 | 2.642E-06 | 12.84 |
| ctg7180001866403_1903_SCT | 19093415 | 3.540E-16 | 35.58 |
| rs864254197 | 19093698 | 2.642E-06 | 12.84 |
| rs864048824 | 19094438 | 3.540E-16 | 35.58 |
| rs863882958 | 19094726 | 5.983E-01 | 0.51 |
| rs864034428 | 19095060 | 6.992E-06 | 11.87 |
| rs864020357 | 19098534 | 9.850E-05 | 9.23 |
| rs863808798 | 19118915 | 4.909E-01 | 0.71 |
| rs863889518 | 19121606 | 3.540E-16 | 35.58 |

| | | | |
|---|---|---|---|
| rs863261265 | 19122801 | 4.909E-01 | 0.71 |
| rs863540518 | 19123299 | 4.696E-03 | 5.36 |
| rs863565552 | 19123704 | 1.828E-02 | 4.00 |
| rs864190920 | 19124146 | 9.169E-01 | 0.09 |
| rs863360143 | 19124890 | 3.540E-16 | 35.58 |
| rs863403978 | 19129310 | 3.540E-16 | 35.58 |
| rs864106897 | 19129402 | 6.115E-03 | 5.10 |
| rs863410946 | 19129698 | 5.374E-01 | 0.62 |
| rs863380259 | 19140354 | 3.540E-16 | 35.58 |
| rs863760013 | 19140509 | 3.540E-16 | 35.58 |
| rs863881957 | 19140703 | 3.441E-01 | 1.07 |
| rs863682040 | 19140768 | 3.441E-01 | 1.07 |
| ctg7180001557189_601_SAG | 19140919 | 6.130E-16 | 35.03 |
| ctg7180001864332_11963_SGT | 19140920 | 6.037E-01 | 0.50 |
| rs863246299 | 19141015 | 2.480E-01 | 1.39 |
| rs863614786 | 19141898 | 4.070E-12 | 26.23 |
| rs863604110 | 19142352 | 4.070E-12 | 26.23 |
| rs863298115 | 19146845 | 3.540E-16 | 35.58 |
| rs863545275 | 19148069 | 3.540E-16 | 35.58 |
| rs863367735 | 19151456 | 3.540E-16 | 35.58 |
| rs864156133 | 19151502 | 3.540E-16 | 35.58 |
| rs863761804 | 19151629 | 3.540E-16 | 35.58 |
| rs863585018 | 19152298 | 3.540E-16 | 35.58 |
| rs863469162 | 19152617 | 3.540E-16 | 35.58 |
| rs864177234 | 19153282 | 3.540E-16 | 35.58 |
| rs863356450 | 19160265 | 3.540E-16 | 35.58 |
| rs864169181 | 19161558 | 3.441E-01 | 1.07 |
| rs863287840 | 19161911 | 3.441E-01 | 1.07 |
| rs863893068 | 19167956 | 8.787E-17 | 36.97 |
| rs863953675 | 19168444 | 3.540E-16 | 35.58 |
| rs863464930 | 19168914 | 2.014E-15 | 33.84 |
| rs863903779 | 19170163 | 3.026E-14 | 31.13 |
| rs864138670 | 19172547 | 1.728E-15 | 33.99 |
| rs864013762 | 19176817 | 3.026E-14 | 31.13 |
| rs864058885 | 19185539 | 1.728E-15 | 33.99 |
| rs864038949 | 19187004 | 7.166E-05 | 9.54 |
| ctg7180001848181_13203_SAG | 19187310 | 2.035E-01 | 1.59 |
| rs863907852 | 19194529 | 1.728E-15 | 33.99 |
| rs863582789 | 19203125 | 5.492E-01 | 0.60 |
| rs863977958 | 19230908 | 2.021E-03 | 6.20 |
| rs863388278 | 19232649 | 1.794E-01 | 1.72 |
| rs863821607 | 19247838 | 3.157E-01 | 1.15 |
| rs864031964 | 19249105 | 1.965E-01 | 1.63 |

| | | | |
|---|---|---|---|
| rs863430709 | 19249788 | 1.965E-01 | 1.63 |
| rs863965270 | 19252011 | 1.965E-01 | 1.63 |
| rs863575174 | 19253089 | 1.965E-01 | 1.63 |
| rs864050148 | 19254654 | 2.069E-03 | 6.18 |
| rs863541276 | 19255154 | 1.965E-01 | 1.63 |
| rs864168424 | 19260022 | 5.198E-04 | 7.56 |
| rs863538804 | 19269705 | 2.291E-09 | 19.89 |
| ctg7180001902815_11658_SAG | 19270046 | 5.492E-01 | 0.60 |
| rs863261312 | 19270596 | 9.020E-10 | 20.83 |
| rs863250331 | 19271060 | 4.054E-09 | 19.32 |
| rs864153919 | 19275782 | 2.246E-08 | 17.61 |
| rs864092962 | 19279098 | 2.045E-01 | 1.59 |
| rs863992350 | 19279255 | 6.377E-07 | 14.27 |
| rs863537347 | 19281062 | 3.241E-01 | 1.13 |
| rs864175148 | 19284305 | 9.448E-01 | 0.06 |
| rs864187608 | 19285401 | 1.017E-08 | 18.40 |
| rs863265930 | 19285976 | 9.939E-04 | 6.91 |
| rs863749747 | 19286936 | 3.437E-03 | 5.67 |
| rs863530042 | 19287078 | 4.565E-03 | 5.39 |
| rs863313687 | 19287560 | 4.565E-03 | 5.39 |
| rs863669620 | 19288417 | 2.497E-03 | 5.99 |
| rs863490286 | 19288985 | 6.340E-10 | 21.18 |
| rs863661159 | 19290159 | 6.606E-05 | 9.62 |
| rs863888145 | 19292315 | 1.091E-10 | 22.94 |
| rs863633141 | 19295827 | 2.613E-02 | 3.64 |
| rs864052472 | 19296862 | 9.888E-11 | 23.04 |
| rs863548467 | 19297779 | 2.005E-01 | 1.61 |
| rs863406273 | 19298897 | 1.198E-01 | 2.12 |
| rs864025048 | 19299502 | 7.002E-06 | 11.87 |
| rs864065881 | 19304485 | 5.399E-03 | 5.22 |
| rs864216474 | 19307972 | 4.935E-01 | 0.71 |
| rs863335535 | 19308404 | 4.935E-01 | 0.71 |
| rs864031125 | 19309772 | 4.935E-01 | 0.71 |
| rs864132037 | 19310772 | 2.302E-06 | 12.98 |
| rs863407349 | 19323334 | 4.638E-03 | 5.37 |
| rs863385810 | 19330797 | 1.311E-02 | 4.33 |
| rs863250308 | 19331271 | 1.640E-02 | 4.11 |
| rs863720013 | 19331308 | 3.013E-05 | 10.41 |
| rs864151096 | 19344468 | 1.311E-02 | 4.33 |
| ctg7180001844909_9843_SAG | 19355435 | 4.068E-03 | 5.50 |
| rs864137208 | 19355971 | 9.453E-01 | 0.06 |
| rs864078587 | 19356038 | 5.880E-10 | 21.25 |
| rs863969489 | 19356072 | 5.880E-10 | 21.25 |

| | | | |
|---|---|---|---|
| rs864140305 | 19361565 | 5.880E-10 | 21.25 |
| rs863437066 | 19366146 | 5.880E-10 | 21.25 |
| rs863345275 | 19366194 | 2.108E-01 | 1.56 |
| rs863360863 | 19369836 | 2.144E-03 | 6.15 |
| rs863448879 | 19371556 | 8.300E-02 | 2.49 |
| rs863576794 | 19371753 | 8.300E-02 | 2.49 |
| rs864210114 | 19371975 | 8.300E-02 | 2.49 |
| rs863842708 | 19372545 | 8.300E-02 | 2.49 |
| rs863547918 | 19372813 | 3.772E-03 | 5.58 |
| rs863961201 | 19374755 | 9.702E-08 | 16.15 |
| rs863588898 | 19375646 | 9.702E-08 | 16.15 |
| rs863360001 | 19376421 | 6.716E-01 | 0.40 |
| rs863818733 | 19377186 | 5.314E-10 | 21.36 |
| rs863751556 | 19377966 | 6.076E-02 | 2.80 |
| rs863936459 | 19391643 | 2.302E-10 | 22.19 |
| rs863789196 | 19402947 | 1.847E-03 | 6.29 |
| rs863234380 | 19403130 | 2.024E-10 | 22.32 |
| rs863346344 | 19406035 | 1.418E-02 | 4.26 |
| ctg7180001215171_1534_SGT | 19406868 | 2.024E-10 | 22.32 |
| rs863772984 | 19407499 | 1.227E-06 | 13.61 |
| rs864171414 | 19414199 | 9.079E-01 | 0.10 |
| rs863977490 | 19414428 | 6.272E-03 | 5.07 |
| rs864117195 | 19415676 | 2.162E-04 | 8.44 |
| rs864092011 | 19418293 | 1.512E-14 | 31.82 |
| rs863338437 | 19427331 | 2.140E-04 | 8.45 |
| rs864097033 | 19428398 | 3.857E-02 | 3.26 |
| rs863241256 | 19441424 | 3.980E-05 | 10.13 |
| rs863708442 | 19446180 | 3.399E-03 | 5.68 |
| rs863578259 | 19446883 | 5.221E-04 | 7.56 |
